# Insight on physicochemical properties governing peptide MS1 response in HPLC-ESI-MS/MS proteomics: A deep learning approach

**DOI:** 10.1101/2023.02.10.527973

**Authors:** Naim Abdul-Khalek, Reinhard Wimmer, Michael Toft Overgaard, Simon Gregersen Echers

## Abstract

Accurate and absolute quantification of individual peptides in complex mixtures is a challenge not easily overcome. A potential solution is the use of quantitative mass spectrometry (MS) based methods, however, current state of the art requires foreground knowledge and isotopically labeled standards for each peptide to be accurately quantified. This increases analytical expenses, time consumption, and labor, limiting the number of peptides that can be quantified. A key step in developing less restrictive label-free quantitative peptidomics methods is understanding of the physicochemical properties of peptides that influence the MS response. In this work, a deep learning model was developed to identify the most relevant physicochemical properties based on repository MS data from equimolar peptide pools. Using an autoencoder with attention mechanism and correlating attention weights with corresponding physicochemical property indices from AAindex1, we were able to obtain insight on the properties governing the peptide-level MS1 response. These properties can be grouped in three main categories related to peptide hydrophobicity, charge, and structural propensities. Moreover, we present a model for predicting the MS1 intensity output based solely on peptide sequence input. Using a refined training dataset, the model predicted log-transformed peptide MS1 intensities with an average error of 11%.

## 1. Introduction

Mass Spectrometry (MS) is a very powerful method for identification and quantification of a wide range of biomolecules present in complex mixtures. The popularity of MS is particularly related to high throughput capability, sensitivity, selectivity, and flexibility, and is therefore utilized in various fields within science and industry for several different applications [1]. The food, pharmaceutical, and medical sectors are just some examples where MS is routinely employed for the identification and quantification of lipids, proteins, peptides, metabolites, and other biomolecular analytes [2–6]. MS is often used in combination with other technologies, particularly chromatography-based methods such as High-Performance Liquid Chromatography (HPLC), to facilitate separation of analytes present in a complex sample prior to MS analysis. Initially, the ionization source is responsible of generating charged analytes in gas phase from an initial solid, liquid or gas phase. Then these charged analytes are discriminated by the mass analyzer based on the mass-to-charge ratio (*m/z*). Tandem mass spectrometry (MS/MS) represents the selective dissociation of intact, well-defined, and charged precursor molecules (i.e. a specific *m/z* from MS1), previously isolated in the mass analyzer, into fragments or ion products. Consequently, MS/MS uncovers more information about the molecular composition and is therefore an excellent platform to identify molecules [7].

The development of soft ionizations techniques such as electrospray ionization (ESI) and matrix-assisted laser desorption/ionization (MALDI) allowed the introduction of MS technology in proteomics research. Although ESI and MALDI represent key breakthroughs in MS-based proteomics, limitations remain due to variability in the ionization efficiency between different peptides and proteins. The variation in the ionization between molecules directly implies that MS technology is not inherently quantitative [8–10]. Through various data normalization approaches, it has been possible to develop methods for label-free, relative quantification with MS [11,12] for different applications. In contrast, absolute quantification by MS requires *a priori* knowledge about the compound(s) to be quantified to develop targeted approaches. Moreover, a standard series or the addition of isotopically labelled reference standards in known concentrations is required to quantify each compound. Thus, absolute quantification methods introduce restrains and limitations to the number of compounds that can be quantified but also higher analytical complexity and especially cost of the MS analysis [13–16]. While efforts have been made towards absolute and label-free quantification on the protein-level [17,18], these approaches still rely on fundamental assumptions regarding the composition of the sample and thus limits the applicable range to protein-level quantification and only for samples of certain origin. Ultimately, there is a need to develop new and universally applicable methods for absolute MS-based quantification on the peptide-level without *a priori* knowledge of the mixture composition, alleviating the large uncertainties associated with using the raw MS1 intensity as a rough pseudo-estimate of peptide abundance [19,20]. To address this challenge, artificial intelligence (AI) is gaining headway for bringing novel solutions to the field of MS-based proteomics [21].

AI is a branch of computer science that focusses on developing systems able to perform tasks which require human like capabilities [22]. At the early stages of AI, programs were developed by explicitly handcrafting an adequately large set of rules to perform specific tasks, in what is known as symbolic AI. But the complexity of certain tasks, particularly when rules cannot be explicitly defined, makes symbolic AI unapplicable. Thus, the concept of machine learning (ML) arose. ML is based on algorithms which generate the rules to perform a specific task by looking at data, without being explicitly programmed. ML works by finding a meaningful transformation of the data, that generates a new and better representation of the data, which is closer to the expected result. Deep learning (DL) is a subfield of ML that, in essence, consists of performing successive transformation of the data across different stages, through a specific arrangement of *layers*, that constitute a particular *deep neural network*, representing the model’s architecture. Therefore, ML and DL are data-centric approaches to develop models to perform specifics tasks. In the field of protein science, ML and DL have facilitated substantial advances for the prediction of e.g., protein structure, protein function, and protein-protein interaction [23–25], but is also rapidly gaining headway in the field of MS-based proteomics. ML- and DL-models have been developed to perform certain tasks such as prediction of peptide retention time, MS/MS fragmentation spectra, and post-translational modifications [21,26,27]. Currently, there are no computational methods to perform absolute peptide quantification based on MS response, since the MS response varies widely between different peptide molecules [21]. In addition to the physicochemical properties of the analytes, in this case peptides, also the experimental setup is bound to influence the results obtained by MS [28]. As such, this task is complex by nature and could therefore potentially be resolved through the application of AI. In recent years, a number of tools have been developed, that exploit DL for prediction of proteotypic peptides, such as AP3 [29], PeptideRanger [30], Typic [31], and AlacatDesigner [32]. Proteotypic peptides are peptides that are well suited for MS analysis as they are released through common sample preparation (i.e., tryptic digest) and are likely to be ionizable and detectable [33]. This makes such peptides optimal choices for e.g., relative quantification between samples in targeted/data-independent analysis or as isotopically labeled surrogates for absolute quantification [34,35]. While such tools may serve as an excellent resource for experimental design in typical proteomics experiments, they focus more on the “what” and not so much on the “why”, when it comes to peptide detectability and ultimately would require custom spike-in to perform absolute quantification. A key in further development towards label-free, absolute quantification on the peptide-level is repurposing of repository data to build sufficiently large datasets suitable for DL [36]. Recent years have also shown substantial progress in this field and efforts towards not only compiling repositories, but also for systematic metadata annotation as well as data extraction and preprocessing [37].

In this study, we investigated the to-date largest repository collection of equimolar peptide MS data [38–40] to develop a DL model capable of providing insight on the physicochemical properties that govern the peptide precursor intensity response (MS1) of peptides in HPLC-ESI-MS/MS analysis - based solely on their amino acid (AA) composition. The AAindex1 (Amino Acid Index Database) is a publicly available collection of 566 indices that describe for instance physicochemical or structural properties and propensities of individual AAs. Each index consists of a set of 20 values that correspond to a specific property of each individual AA [41]. The results obtained in this study provide a better fundamental understanding of the behavior of peptides within the mass spectrometer. Moreover, a model was developed to predict peptide MS1 response as a function of the AA composition. The presented work is of great relevance for development of more advanced models to predict e.g., peptide detectability and to facilitate advances in label-free, absolute peptide quantification. Furthermore, the improved understanding may aid future improvements and guide design of new MS technologies to bypass current limitations.

## 2. Results and Discussion

The models created for this work are autoencoders with attention mechanism, where the encoder transforms the input data into a more meaningful representation. Subsequently, the attention mechanism assigns weights of contribution to each of the elements of the sequence. Ultimately, the decoder uses the information of the encoded sequences and the attention weights to predict the log-transformed MS1 intensity output. To ensure satisfactory performance of the fundamental architecture, the model was initially developed using artificial datasets with known ground truth.

### 2.1. Model development using artificial datasets

Proof-of-concept models were initially trained to explore their efficacy, capacity, and limitations for the model itself. For this purpose, artificial datasets were generated consisting of numeric sequences, where each sequence had a scalar value assigned, emulating an experimental “intensity output value” (Table S1). Fixed contributions were assigned to each unique element of the input sequences and different sets of contributions as well as sequence formats were analyzed for the proof-of-concept models. The first dataset format consisted of sequences with a length of 4 to 8 elements comprised of 9 unique numbers (from 1 to 9). Two sets of contribution values were assigned to each unique elements (Table S2) to test the architecture. One set of contributions consisted in an increasingly linear contributions of each unique elements with values ranging from 10 to 90, obtaining mean absolute percentage error (MAPE) of 0.79% and Pearson correlation coefficient (PCC) of 0.999 between the attention weights and the contribution values (Table S3). The second set of contributions consisted of a linear contribution ranging from 1 to 9 for odd numbers and a non-linear contribution ranging from 10 to 160 for even numbers (Table S2). This was done to increase the dynamic range of contributions as well as the gap of contributions between the unique elements. The second set of contributions resulted in a MAPE of 0.56% and PCC of 0.980 (Table S3). In both cases, the attention mechanism successfully identified the relevancy of individual element (Fig. S1) with excellent correlation between real and predicted values (Fig. S2).

The second artificial dataset format consisted of sequences with a length of 7 to 40 elements comprised by 20 unique numbers (from 1 to 20). Similarly, two sets of contribution values were assigned to each unique element (Table S4) to test the architecture. The second dataset format is meant to better emulate the characteristics of the real data, namely the length of the sequences, the order of magnitude of the MS intensity output, and the number of unique elements (20 AA). One set of contributions consisted of an increasingly linear contribution of each unique element with values ranging from 1.0×10^7^ to 2.0×10^8^, obtaining MAPE of 0.60% and PCC of 0.999 (Table S5). The second set consisted of four groups of elements having different orders of magnitude of contributions ranging from 0 to 2.0×10^9^ with a MAPE of 3.15% and PCC of 0.981 (Table S5). Similarly, to the first dataset format, the attention mechanism successfully identified relevant contributions (Fig. S3) and the models obtained strong correlation between predicted and real values (Fig. S4). For all artificial datasets a log-transformation was applied to the output values, which is a common transformation when dealing with non-linear data representing a large dynamic range of values [42–44]. For proof-of-concept models 3 and 4, log-transformations were found a prerequisite to find a solution for the task, while it did not seem to have an impact on the performance of proof-of-concept models 1 and 2.

In general, the models were able to assign the correct contribution of each of the unique elements in the different sequence formats. The lowest correlation was observed with the most complex sequence (20 unique elements) when the fixed contribution values had a very large dynamic range with a maximum contribution of 2×10^9^ and a minimum of 0 (Proof-of-concept model 4). This particular model was unable to assign the correct average contribution of the unique sequence elements which had a contribution lower than 2×10^6^ (3 orders of magnitude below the maximum contribution), although it was able to group elements by order of magnitude of their contribution (Table S6, Fig. S3B). Nevertheless, all the models were capable of determining the most relevant contributing elements of the sequences in a satisfactory manner and make highly accurate prediction of the output value, thereby validating applicability of the architecture. Moreover, accurate predictions were obtained in scenarios accounting for a uniform distribution of the contribution by each element, but also scenarios accounting for high contrast between contributions of individual elements.

### 2.2. Dataset clean-up and initial model implementation

After the model architecture was proven effective using artificial datasets, the next step was to train and test the model with real MS data. For this purpose, we extracted the MaxQuant [45] output datafiles from the PROSIT and ProteomicsDB datasets (PRIDE identifiers PXD004732 [38], PXD010595 [39], and PXD021013 [40]), that were produced by orbitrap analysis of synthetic, equimolar peptide pools. To ensure optimal training of DL models using real datasets, these should be of sufficient quality. Therefore, we initially inspected the datasets with the aim of investigating variability and data quality. The cumulative database consists of 4,016,044 peptide identifications representing 1,331,904 unique peptides. As commonly applied in proteomics studies, reverse sequences were eliminated as false positives and potential contaminants removed to improve data reliability. Because the majority of peptides (865,325 or 64.97%) were analyzed and identified in more than one pool, this allowed us to investigate the variability of the MS1 intensity data by computing the coefficient of variation (CV) for the different intensity measurements of the same peptides (Fig. 1A). MS1 intensities show a high variability with CVs exceeding 400% for some peptides. Thus, the CV was used to filter peptides with high variability (CV > 30%) from the initial dataset. In MaxQuant output data, there are additional metrics, which are commonly employed for downstream filtering and processing prior to further analysis. Some metrics relate to quality of identification, as for instance the posterior error probability (PEP). While PEP is used in the calculation of the peptide/protein score by the MaxQuant built-in search engine Andromeda [46], other factors are also accounted for when calculating this score [47]. The score is directly applied in the filtering during initial MaxQuant analysis through the false discovery rate (FDR), assigned by the user. Consequently, the PEP may also be used as a stand-alone metric to perform further quality-based filtering. As such, we used PEP as a parameter to remove peptides that might have been incorrectly identified by defining a maximum threshold of 1% (i.e., removing peptides with PEP > 0.01).

**Figure 1.**
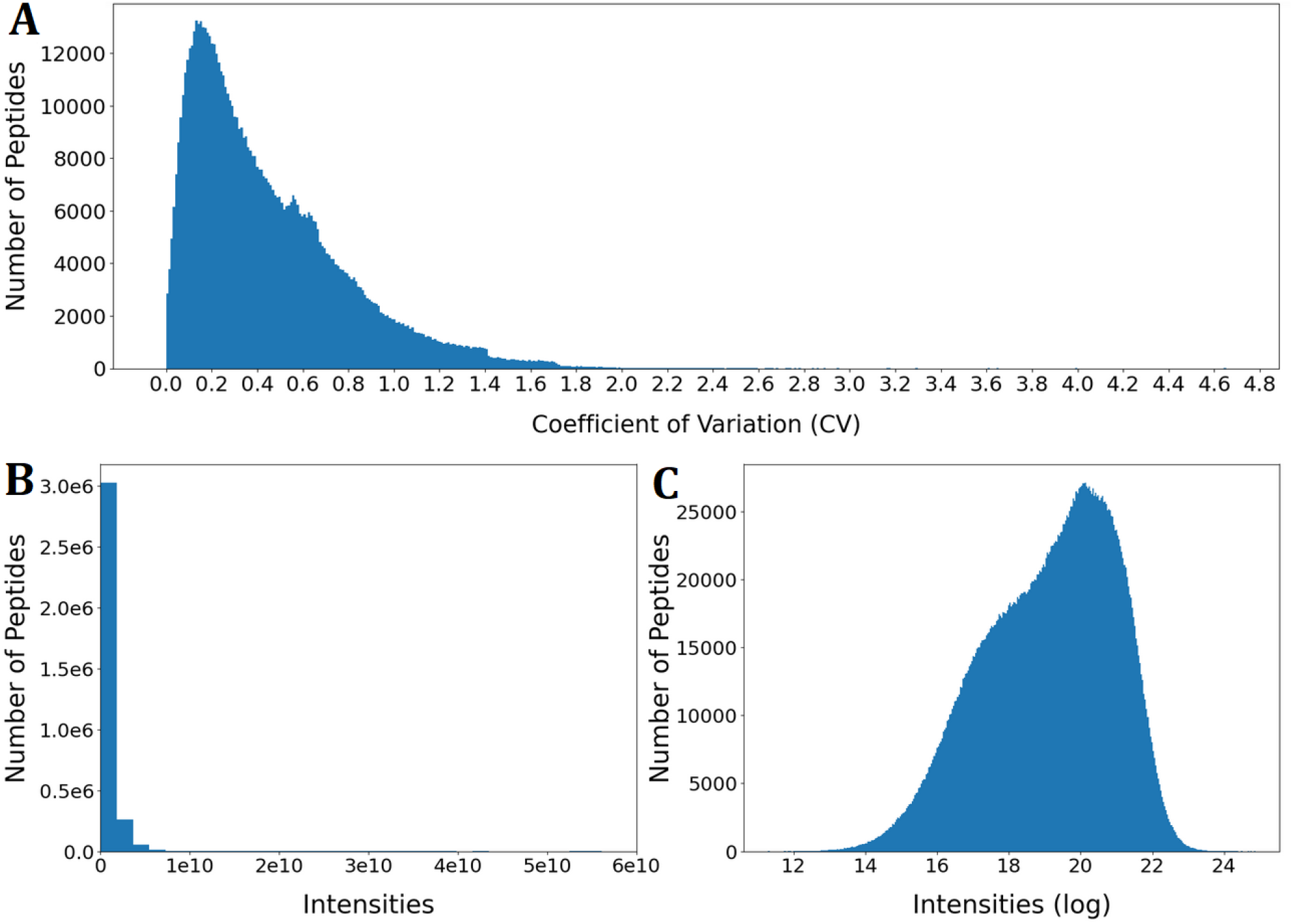
Initial exploration of the cumulated dataset. A) Distribution of coefficient of variation (CV) for peptides with more than one measurement (Bin size ≈ 0.010). B) Distribution for raw MS1 intensities for filtered data (Bin size ≈ 1.8×109. C) Distribution of log-transformed MS1 intensity data for filtered data (Bin size ≈ 0.036).

The final dataset consisted of 320.741 unique peptides. Once the data was filtered, the MS1 intensity output were log-transformed with the natural logarithm. This allowed to reduce the dynamic range of intensity outputs, reducing the impact of high-intensity peaks, and generating a distribution closer to normal (Fig. 1B and 1C). The overall effect of filtering was primarily reducing the size of the dataset without affecting the distribution or dynamic range substantially (Fig. S5).

During initial model training and optimization, two consistent patterns were observed in the attention weights for each of the 20 AAs. While the patterns are quite different, the performance of the different models were comparable, with MAPEs generally in the range from 12-18% (see supplementary information). The first pattern frequently highlighted the influence of bulky hydrophobic (i.e., leucine (Leu), isoleucine (Ile), and valine (Val)) and aromatic (i.e., tryptophan (Trp), phenylalanine (Phe), and tyrosine (Tyr)) AAs. In contrast, the second pattern primarily highlighted a high contribution by positively charged AAs (i.e., arginine (Arg) and lysine (Lys), and to a lesser extend histidine (His)). Although these patterns were frequently observed during the training and optimization process, the exact results, namely the attention weights and their distribution, were not consistently reproducible due to the stochastic nature of the algorithm. In other words, the order of the AAs occasionally shuffled, but the overall pattern remained intact. After computing the correlation between the attention weights and the parameters contained in AAindex1, certain physicochemical properties were reproducibly identified for the highly contributing AAs within the two different patterns emerging. To illustrate this, representative models were selected for further analysis.

### 2.3. Representative Model 1: Bulky Hydrophobic and Aromatic Amino Acids

For the first representative model, the highest attention weights are given mainly to bulky hydrophobic and aromatic AAs. While generally considered hydrophobic, alanine (Ala) shows a substantially lower contribution, which may likely be related to the lower mass and/or volume compared to other hydrophobic AAs. Interestingly, sulphur-containing AAs (i.e., cysteine (Cys) and methionine (Met) also receive some attention from the model. While Cys is not considered bulky, the sulphur has been alkylated (carbamidomethyl), increasing the size of the side chain substantially. The attention weights are, however, modest (Fig. 2, Table S6), which might relate to the inclusion of sulphur (and terminal amide in the case of caramidomethylcysteine), increasing the overall polarity of the side chain compared to aliphatic AAs. Similarly, Tyr receives lower attention compared to the other aromatic AAs (Trp and Phe), which may be related to the hydroxyl group increasing side chain polarity. While Phe is generally considered more hydrophobic, Trp receives the highest attention of all AAs. While this may relate to the larger size of the side chain, Trp is also known to function as a gas-phase charge stabilizer through the indole moiety [48,49], which could improve stability of the precursor ion, adding to the overall influence on the MS1 response.

**Figure 2.**
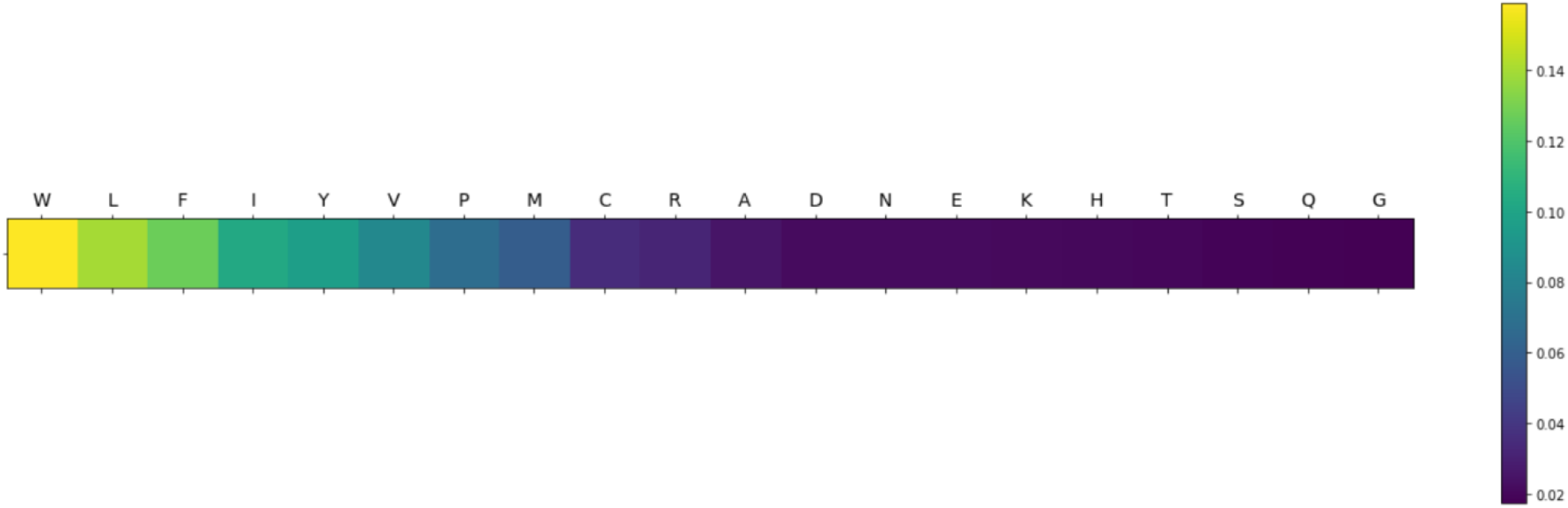
Graphical representation of the attention weights of each AA for representative model 1. The color coding indicates the assigned contribution of each AA to the prediction of the MS1 intensity output from high contribution (yellow) gradually decreasing to low contribution (dark blue).

Computing the correlation between attention weights and AAindex1, parameters related with hydrophobicity (Table 1, Table S7) were found of great relevance. This indicates a strong correlation between hydrophobicity and the MS1 intensity measurement. *Retention coefficients and hydrophobicity indices* were often found relevant. In reverse phase (RP) chromatography with applied solvent gradients going from high towards low polarity, higher peptide retention times are a result of higher peptide hydrophobicity. As acetonitrile, which is commonly used as the organic phase in LC-MS/MS-based proteomics, has a higher vapor pressure than water, it is substantially more volatile. Thus, when peptides with higher retention times (i.e., eluting late) reach the ion source, the solvent is easier to evaporate. Moreover, hydrophobic peptides are generally more inclined to be in the organic phase [50], explaining why *partition coefficient* is another relevant property identified. Furthermore, hydrophobic peptides are usually located towards the surface of the droplets [51,52], which is also reflected by the identification of different *transfer free energy* properties as relevant. These factors illustrate why more hydrophobic peptides might generally have a better ionization efficiency in gradient RP-HPLC. Other studies have found a direct empirical correlation between ionization efficiency and retention times, corroborating these findings [50,52,53].

**Table 1.**
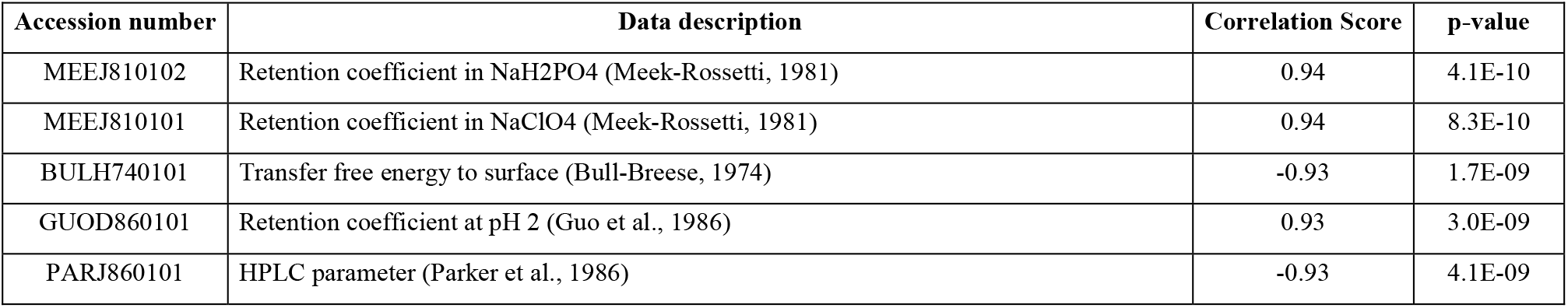
Top 5 relevant physicochemical properties from the AAindex1 identified by correlating the indices with the attention weights of representative model 1.

Other computational approaches have found results similar to our findings [29,54–57]. Jarnuczak *et al*. (2016) found that in complex mixtures, there is a weak non-linear relationship between ionization efficiency and hydrophobicity, which they argue might be linear in a simpler mixture [58]. The authors also show that ionization efficiency is hampered at very low and high organic concentration of the mobile phase, as “weak flyers” were observed at both low and high organic concentration of the mobile phase. They state that at very high organic concentrations, there is an increased basicity in acetonitrile within the gas phase, which interferes with the ionization of peptides. Taken together, this presents substantial evidence of the influence of peptide hydrophobicity on ionization efficiency and thus MS1 response in RP-HPLC-ESI-MS/MS.

As the hydrophobicity and retention coefficients were determined to be highly relevant for peptide response, we investigated if this was directly reflected in the filtered dataset. While two indices showed higher correlations with the attention weights from representative model 1 (Table 1), these are retention coefficient in solvents not common employed in ESI-MS. Consequently, we computed the next three indices (BULH740101, GUOD860101, and PARJ860101) for all peptides as both sum and mean and plotted against the peptide MS1 intensity (Fig. S6). There does not seem to be any direct correlation and thus, intensity response cannot be predicted based solely on hydrophobicity. While higher responses are observed in certain ranges for the different metrics, these merely represent a higher density of datapoints. Nevertheless, the model identifies hydrophobicity as relevant, but the property is not descriptive as a stand-alone variable, and hence the model is finding more complex patterns within the data.

### 2.4. Representative Model 2: Positively Charged AAs

In the second commonly observed pattern, high relevance of AAs with positively charge side chains (Arg, Lys, and to a lesser extent, His) was observed (Fig. 3, Table S8). Correlating attention weights with AAindex1, parameters related with peptide charge were, not surprisingly, found to be very important for this model. *Positive charge* and *net charge* had PCCs of 0.93 and 0.74 with p-values of < 2E-09 and < 2E-04, respectively (Table S9). Since the samples were originally analyzed in positive mode ESI-MS using an acidified solvent (0.1% formic acid), that makes *positive charge* a very intuitive property. The parameter precisely points to those AAs that most likely will be positively charged due to side chain protonation at acidic pH. Thus, the presence of Arg, Lys, and His in a peptide most likely will increase the probability of getting a positively charged ion during ionization. Other studies have also found this particular property of high relevance [55,58,59].

**Figure 3.**
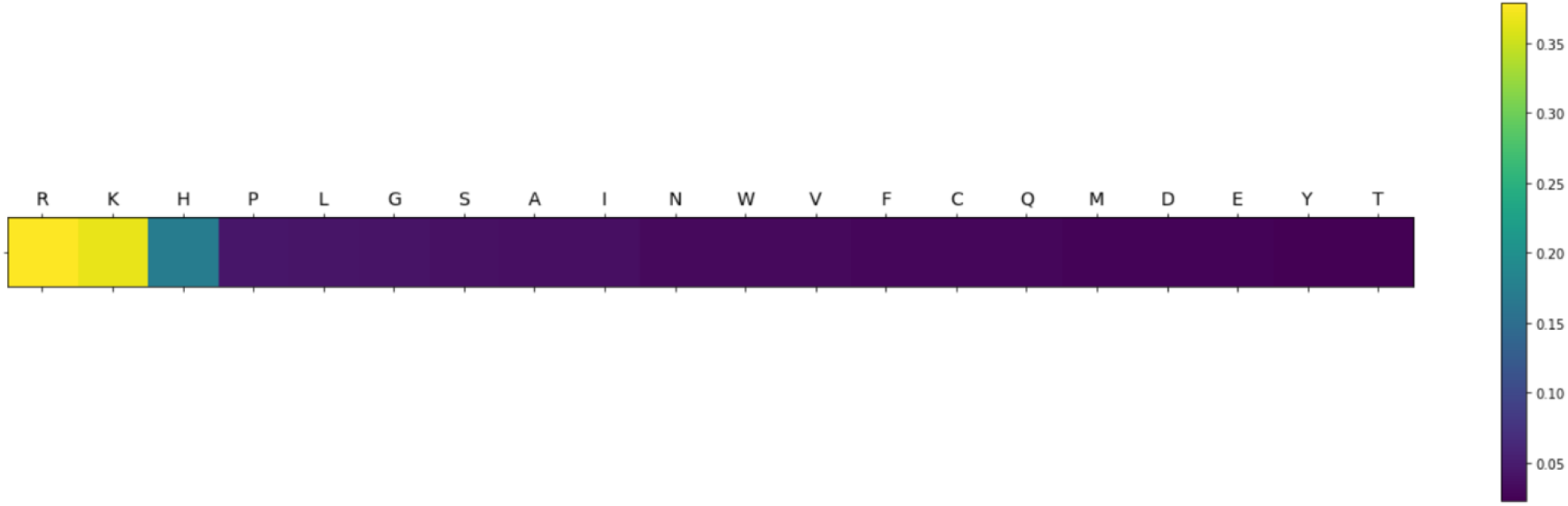
Graphical representation of the attention weights of each AA for representative model 2. The color coding indicates the assigned contribution of each AA to the prediction of the MS1 intensity output from high contribution (yellow) gradually decreasing to low contribution (dark blue).

To investigate if the relevance of positive charge was directly reflected in the filtered dataset, the number of positively charged AAs (Arg, Lys, and His), net charge at pH 7, and net charge at pH 3 (reflecting the acidic environment used during positive mode ESI-MS) was determined for individual peptides and plotted against MS1 intensity (Figure S7A-D). Moreover, these charge-related metrics were also determined in a length-normalized version (charge/length) to investigate the interplay between the two physicochemical properties (Fig. S7E-F)). As found for hydrophobicity descriptors in relation to representative model 1, there is no direct correlation between charge and MS1 intensity, also indicating a more complex interplay between different variables, which the model is able to identify. We also investigated different combinations of hydrophobicity indices and charge (i.e., ratios and products), but also here found these metrics insufficient to describe MS1 intensity (data not shown).

#### 2.4.1. Sub-distributions and Search Parameter-based Data Segmentation

The distribution of the log transformed MS1 intensities in the filtered dataset (Fig. 1C) is to no extent normally distributed and seems to potentially contain more than one distribution. To investigate this, the dataset was segmented according to variable parameters in the MaxQuant metadata related to specified enzymatic digestion and search parameters. When grouping the data based on the specified enzyme and the enzyme mode (i.e., specific, semi-specific, or unspecific *in silico* digestion) [38–40] applied for processing the raw data (Fig. 4), the presence of sub-distributions become evident.

**Figure 4.**
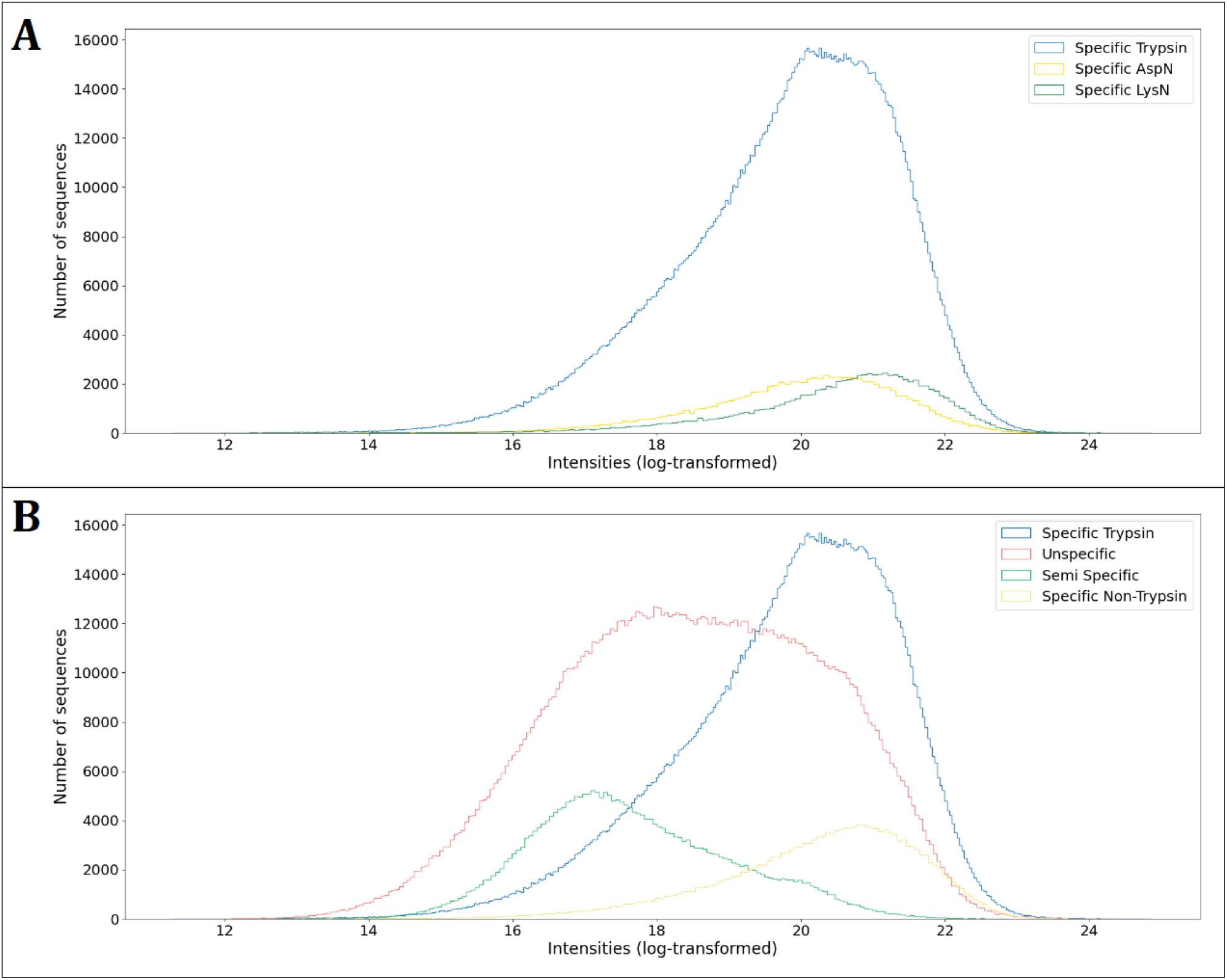
Histograms of log-transformed peptide MS1 intensity outputs by “Enzyme” and “Enzyme Mode”. A) Histogram of peptides quantified using “specific digestion”. B) Histogram with peptides grouped by MaxQuant “enzyme mode” and distinguishing between tryptic or non-tryptic peptides using “specific digestion”.

Peptides searched with a specific enzyme digestion (Trypsin, LysN, and AspN) have a higher median value of intensity than peptides search with unspecific or semi-specific enzymes (Fig. 4A-B, Table 2). Trypsin generates peptides with a C-terminus constituted by Arg or Lys, LysN produces peptides with a N-terminal Lys, while AspN releases peptides with an N-terminal aspartic acid (Asp). While all these specific terminal AAs have electrically charged side chains, Arg and Lys are positively charged while Asp is negatively charged. Distribution of log transformed MS1 intensities seem to suggest that charged AAs, especially when located at the peptide termini, may have a direct effect on the intensity output in MS1. Interestingly, Asp was not found of high relevance (Fig. 3). Asp will not be charged under acidic pH used in positive mode ESI-MS, and therefore constitute an unchanged, polar residue. As such, the higher median intensity of his subset may reflect a potential proximity effect of carboxylic acid moiety and the N-terminal charged amine but may also represent that the peptide composition in the subset is indigenously more suitable for MS detection and hence provides higher MS1 response.

**Table 2.**
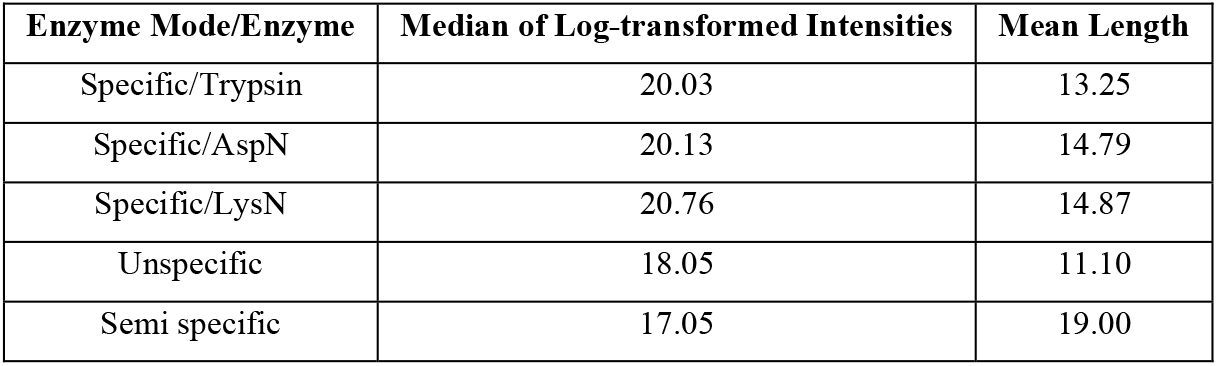
Median values of log-transformed intensities and mean length of peptides (whole dataset) grouped by “Enzyme Mode” and “Enzyme” specified in MaxQuant metadata.

| Enzyme Mode/Enzyme | Median of Log-transformed Intensities | Mean Length |
| --- | --- | --- |
| Specific/Trypsin | 20.03 | 13.25 |
| Specific/AspN | 20.13 | 14.79 |
| Specific/LysN | 20.76 | 14.87 |
| Unspecific | 18.05 | 11.10 |
| Semi specific | 17.05 | 19.00 |

It is important to highlight that peptides searched with semi-specific setting may have shown lower intensity values (Fig. 4B) for two reasons: Firstly, the pools used for these analyses contained longer peptide (average > 25 AAs [40]) than in the other datasets, which can generate a reduction of the intensity measurement due to a bias against longer peptides in the orbitrap mass analyzer [58,60–62]. Secondly, the search mode allowed to find full-length synthetic peptides as well as truncation obtained as incomplete synthesis products [40]. As the pool equimolarity correspond to the full-length peptide, the abundances of truncated forms are expected in substantially lower abundance, thus reducing the intensity values of the truncated sequences detected (Fig. S8C-D). This directly compromises the equimolar prerequisite for the sequence-centric analysis performed in this study and ultimately introduce bias and reduced reliability of the dataset. This becomes particularly evident through the mean length of the identified peptides using semi-specific search (Table 2), as this (19 AAs) is substantially lower than the specified average length for the peptide pools (> 25 AAs). For unspecific *in silico* digestion (Fig. 4B), there does not clearly seem to be higher response for peptides with a C-terminal Arg or Lys, although tryptic peptides identified in unspecific searches do represent the high responders too (Fig. S9-A). The apparent bimodal distribution indicates that additional properties account for the segregation of this subset in to (at least) two additional subsets. Interestingly, there seems to be a more consistent increase in MS1 intensity for peptides containing Arg or Lys (anywhere in the sequence) in comparison with those that do not (Fig. S9-B). This observation further substantiates the importance of positively charged AAs for a high ionization efficiency and thus MS1 response (corroborating the findings from representative model 2), while length itself does not seem to correlate directly with MS1 response in general terms (Figure S8-A,C).While length does seem to influence response to some degree, this may simply be related to the fact that these lengths are in general overrepresented in the dataset (Fig. S8-B, D)

### 2.5. The influence of structural properties

In addition to hydrophobicity and charge, a number of physicochemical properties are identified as relevant in the two representative models, which relate to protein and peptide structural properties (Tables S7 and S9). Although many of these properties are also related to e.g., hydrophobicity, they also contain information on structural aspects, as these are often related. For instance, one of the properties showing high correlation with attention weights is the *Atom-based hydrophobic moment*. This parameter quantifies the strength of the periodicity in the polar or hydrophobic nature of the constituent amino acids of a sequence, which is related to the stability and type of structure as well as its functions [63]. Other properties such as *Entropy of formation, solvation free energy*, and *Weights from the IFH scale* are also found relevant. These properties are related with the thermodynamics of protein and peptide conformation and stability [64–66]. *Energy transfer from out to in(95%buried*) and *Buriability* “provides a quantitative measure of the driving force for the burial of a residue”, thereby describing polarity-driven, tertiary conformational properties [67]. While the *Isoelectric point* is a parameter related to charge, it also describes electrostatic interactions between AA side chains, which affect protein and peptide structure [68,69].

There are, however, also important properties that are more directly related to structural aspects of peptides and proteins. For instance, the *Helix termination parameter at position j-2,j-1,j* refers to the formation probability of secondary structures, here specifically α-helices, in peptides [69]. Peptides and proteins can form secondary and tertiary structures not only in solution, but also in the gas phase [70–72]. Studies have shown that peptides with stable α-helical and β-sheet structures in solution have lower intensity response than corresponding structurally disturbed analog peptides (L- to D-AA substitution) in MALDI-MS [73]. This indicates that peptide solution-phase structure could have a significant influence on the MS1 response. Moreover, it has been observed that the fragmentation of protonated peptides is influenced by the peptide’s gas-phase secondary structure and in particular acid-base interactions and charge solvation in the gas phase [74], corroborating that proximity-based intramolecular interactions are indeed of importance for precursor stability during MS analysis. Consequently, the identification of a peptide is influenced by both the peptide primary structure and the consequential secondary structure in the gas phase. In-source fragmentation would lead not only to a lower MS1 response, but also a reduced proportion of the precursor peptide available for MS/MS identification. Furthermore, studies using MALDI-MS have shown that the conformation of peptides in the gas-phase is not necessarily the same than in solution-phase [75]. While ionization method in these studies differs from ESI considered here, the phase transition is still highly relevant and considered of importance in relation to ionization efficiency and thus peptide MS1 response in ESI-MS/MS. Moreover, other studies with computational approaches have similarly found structural properties of significant relevance for MS analysis [29,54,55,57,58]. Based on these findings, peptide structure appears a key factor affecting the MS1 response and an important source of variability in intensity measurements.

### 2.6. Model Performance Optimization and Sequence-based Intensity Prediction

The presented models were evaluated with the test datasets and their performances were expressed through MAPE, showing the percentual distance between the real and predicted MS output intensities. The proof-of-concept models displayed a MAPE between 0.56 and 3.2 % (Tables S3 and S5) with an almost perfect correlation between expected and predicted values (Figs. S2 and S4). This shows that the models have an exceptional performance with the artificial data, not only identifying the average contribution of each unique elements of the sequence but also predicting the expected output. Using the repository MS data, initially all the filtered data was used to train and test the models, obtaining an average MAPE of 14.9% (Supplementary Table S10). Nevertheless, there are substantial differences in intensity distributions based on the applied enzyme and enzyme mode setting used during the data search (Fig. 4B). Therefore, to improve the model performances the model was trained and tested only using a specific subset of the data, namely the Specific/Tryptic peptides, as the remaining subsets had substantial uncertainties, as previously discussed.

When doing this, the MAPE was reduced to 10.5% (Table S10), resulting in a relative reduction in the error of 30% for log-transformed intensities but also an impressive 57% for raw intensities (MAPE=85%) compared to the average MAPEs for the two representative models (average MAPE=198%). The model obtained a PCC of 0.68 between log-transformed intensities and predictions and a PCC of 0.64 between raw intensities and predicted values (Fig. 5B-C). The attention weights of the model consistently focus on the bulkier hydrophobic and aromatic AAs (Fig. 5A and Table S11), thereby showing similar attention patterns as representative model 1 on the whole filtered dataset, and thus giving higher relevance to the hydrophobicity-related properties (Table S12). This might indicate that the transition of peptides from liquid to gas phase and charge stabilization are key factors in sequence-based variability for MS1 intensity measurements. Moreover, this may also simply indicate that the model is now focusing on hydrophobicity because all the peptides in the particular data subset are tryptic. As all peptides feature a C-terminal Arg/Lys, charge may not be of descriptive relevance. This could further indicate that representative model 2 in fact focused more on identification of the specific/tryptic subset, as these peptides overall show a higher MS1 intensity compared to the semi-specific and unspecific subsets (Table 2). To investigate this further, we determined charge-related metrics for these subset peptides and investigated the correlation with the intensity outputs (Fig. S10). Here we see that neither charge nor number of positively charged AAs seem to in any way be descriptive of MS1 intensity variation between peptides, as observed in the previous models. Moreover, the relationship between MAPE (%) of each prediction and peptide length was investigated showing no correlation, indicating that the model has no bias regarding peptide length (Fig. S11).

**Figure 5.**
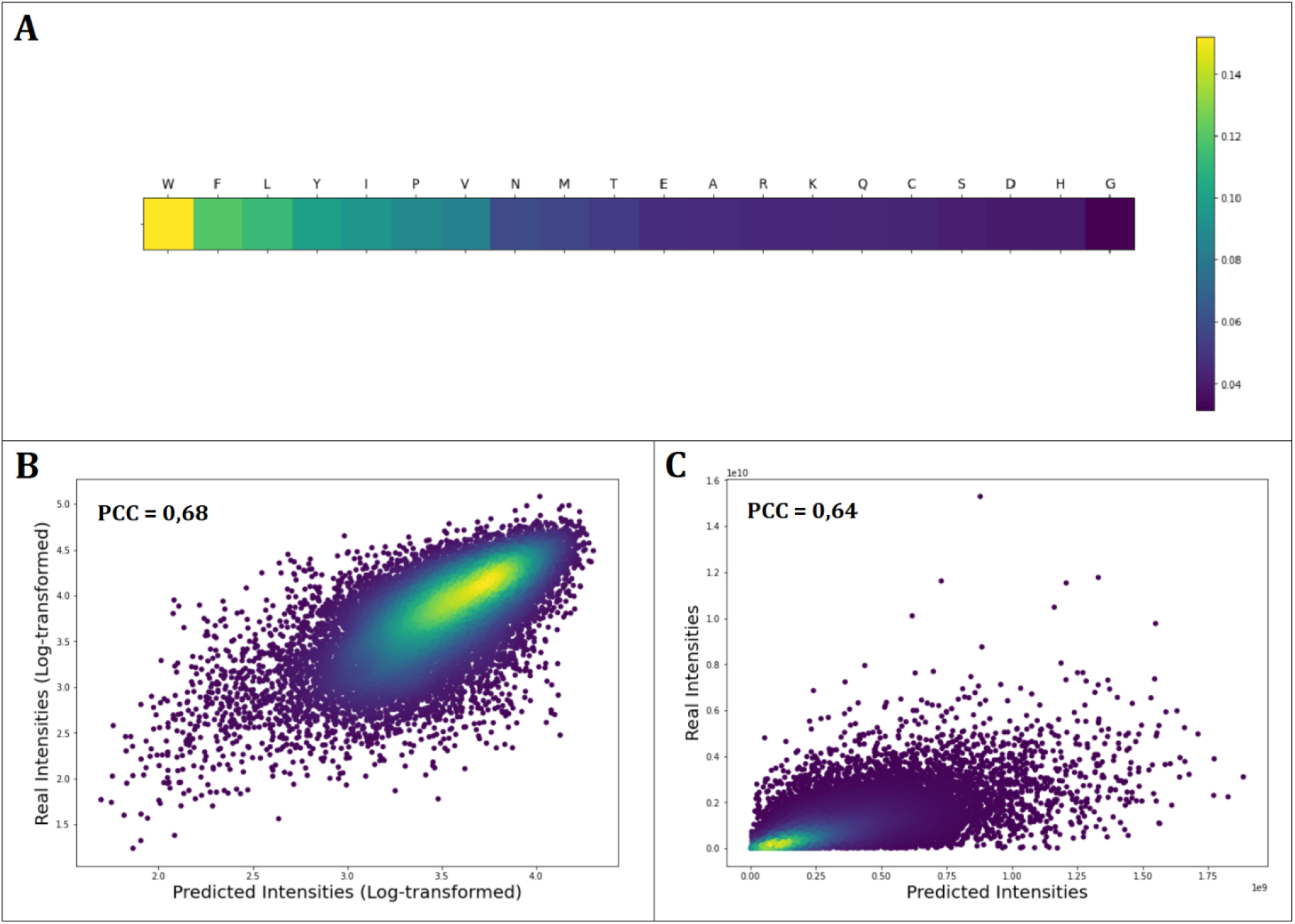
Attention weights and model performance results for the final mode (Specific/Tryptic peptides only). A) Graphical representation of the attention weights of individual AAs. The color coding indicates the assigned contribution of each AA to the MS1 intensity output given by the model. B) Scatter and density plot of the real log-transformed intensities vs the predicted log-transformed intensities. C) Scatter and density plot of the real intensities vs the predicted intensities.

When evaluating model performance, it is important to take into consideration that the models were trained only providing the sequence information and the corresponding MS1 intensity output without explicitly defining any physicochemical properties to be important. Nevertheless, the models seem to be able to identify properties by themselves, that are aligned with previous empirical studies. Furthermore, as there is a clear correlation between predicted and real MS1 intensity outputs (both raw and log-transformed), this shows that the models are effectively extracting meaningful information from the peptide sequences to predict intensity. Certainly, there is a limit to how much the information from the sequences can explain the MS1 intensity output, since there are other sources of variability outside the MS instrumentation. Those limitation arise from sources such reproducibility in sampling and sample preparation [77–79], the type of MS technology employed [1,58,79–82], as well as the raw data processing [83,84]. Moreover, it is essential to remember that there is high variability in the MS intensity output for the same peptides in this particular dataset. Consequently, building more robust datasets and designing standardized experimental protocols that allows to generate more consistent measurement are key factors to building models that can accurately predict peptide intensities and account for intrinsic variability. Such models can ultimately be applied to estimate absolute peptide quantification without the need of isotopically labeled surrogate peptides.

## 3. Conclusions

In this study, a deep learning neural network with attention mechanism was used to determine the relevance of each of the 20 natural amino acids on the MS1 signal response from peptides in HPLC-ESI-MS/MS analysis of equimolar peptide pools. The initial models were capable of predicting log-transformed peptide intensity with an average MAPE of 14.9%. The attention weights from the models were correlated with the physicochemical property indices contained in the AAindex1 to identify which physicochemical properties play an important role in the behavior of peptides in MS, as well as their impact on MS1 intensity measurements. Hydrophobicity, charge, and structure were identified as important relevant properties governing the peptide MS1 responses. These parameters were not directly reflected in the data, but extractable using AI. Following further segmentation of the dataset, the model was trained on only specific/tryptic peptides, thereby improving the model performance, and reducing MAPE for log intensity prediction to 10.5%. The model performance is likely to be improved by generating more accurate and robust datasets as well as experimental protocols to normalize between individual MS runs. Overall, the information generated in this study is of great relevance to understand the key factors influencing the results obtained in HPLC-ESI-MS/MS peptide analysis. This information can also be used to build more advanced models for peptide detectability and peptide quantification, as well as to enhance or even design new MS technology to overcome current limitations.

## 4. Materials and Methods

### 4.1. Data

Artificial datasets were generated for proof of concept of the deep neural network model. Each dataset contained 100.000 training sequences from which 20% were randomly selected as validation dataset. An additional 10.000 sequences were generated as test dataset. The first dataset format consisted of arrays of random numbers from 1 to 9 with a variable length of 4 to 8 elements. The second dataset format contained sequences with random numbers from 1 to 20 with a variable length of 7 to 40, to better emulate the structure of the real peptide MS datasets. Each number was assigned a fixed value or contribution and the contribution of all the elements in each sequence were added to obtain the output value of the corresponding sequence (see supplementary material).

The experiment data used in this study was collected from the PRIDE repository with the identifiers PXD004732 [38], PXD010595 [39], and PXD021013 [40]. These databases were built by analyzing pools of approximately 1000 synthetic peptides with equimolar concentrations. RAW data was analyzed using specific, semi-specific, and unspecific enzymes settings [45] in their respective studies, using Trypsin, LysN, and AspN as specified proteases. In all studies, peptide pools were subjected to liquid chromatography using a Dionex 3000 HPLC system (Thermo Fisher Scientific) coupled online to an Orbitrap Fusion Lumos mass spectrometer (Thermo Fisher Scientific) [38–40]. This ensured experimental comparability between studies and was a prerequisite for inclusion in the cumulated database employed in this study. From the peptide-level MaxQuant [45] output files (peptides.txt, summary.txt) and Sample and Data Relationship File (SDRF), several data features were extracted using a custom Python script. Each pool analyzed had a corresponding zip file containing the peptide.txt and summary.txt files. The final results of each analysis were extracted from the peptide.txt file (sequence of identified peptides, MS1 intensities, PEP scores, etc.) while the enzyme and enzyme mode settings were extracted from the summary.txt files. The SDRF files contains information relating each pool zip file with its specific experimental setups. A unique CSV files was generated for each pool unifying the information in the previously mentioned files, which later were merged into a single CSV file comprising all the information of the repositories PXD004732, PXD010595 and PXD021013.

### 4.2. Data filtering and pre-processing

To build the best possible model, the data (4,016,044 identified peptides) was initially filtered with the intention of reducing noise, thereby improving data quality for the training and testing process of the models. The artificial data did not require filtering. The data was initially filtered using quality-based criteria:

All peptide sequences with a PEP score equal to or higher than 0.01were removed (587,374 peptides).
Reverse sequences were excluded (414 peptides).
Peptides determined as potential contaminants were not considered (22,257 peptides).
Peptides with intensity measurement equal to zero were discarded (40,798 peptides).

Following initial filtering, the dataset was further processed and filtered suing replication- and variation-based criteria

- Peptide replicates were merged (2,316,063 peptides). For each peptide, the median intensity was used for the analysis.
- Peptides with intensity values with a coefficient of variation (CV) higher than 0.3 (Eq. 1) were dismissed (728,397 peptides).

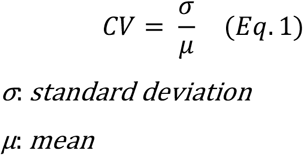

Ultimately, only peptides with more than one measurement were included in the filtered dataset, which consisted of 320,741 unique peptide entries.

To improve the model’s performance the initial filtering was not only followed by reduction of variation and removal of replicates, but started by subsetting the datasets for those peptides searched and identified with MaxQuant using Specific enzyme setting, and Trypsin as protease:

- Peptides that were not searched with Specific Enzyme mode were removed (1,598,623 peptides).
- Non-tryptic peptides were discarded (184,918 peptides)
- Replicate peptide measurements were merged (1,177,976 peptides). For each peptide, the median intensity was used for the analysis.
- Peptides with intensity values with a coefficient of variation higher than 0.3 (Eq. 1) were dismissed (224,462 peptides).

Therefore, only peptides with more than one measurement will be considered. The final number of peptides is 179,222.

Following filtering, peptide intensity values, which are continuous values, were log transformed (natural logarithm) since the data shows an exponential behavior with very high intensities values over a large dynamic range. The log transformation allows to generate a distribution closer to normal and then the intensity values were scaled between a specific range of values, using the *MinMaxScaler* function from Scikit-learn library [85] which was optimized (the same was done to artificial data) by trying different ranges to improve the models’ performances. Log-transformation of intensity data has been previously used to process MS data [43,44,86–88]. Ultimately, data was randomly split into a training (72%), validation (18%), and test sets (10%).

### 4.3. Model Architecture

The model architecture is an autoencoder with attention mechanism. The encoder consists of one bi-directional recurrent neural network layer with Gated Recurrent Units (GRU) [89] and an attention mechanism [90]. The decoder has the same configuration as the encoder (Fig. 6) in addition to a dense layer with one unit corresponding to the predicted intensity. The recurrent layers and the latent space have the same dimensions, while a different number of units and batch sizes were used during training and optimization (Tables S3, S5 and S10).

**Figure 6.**
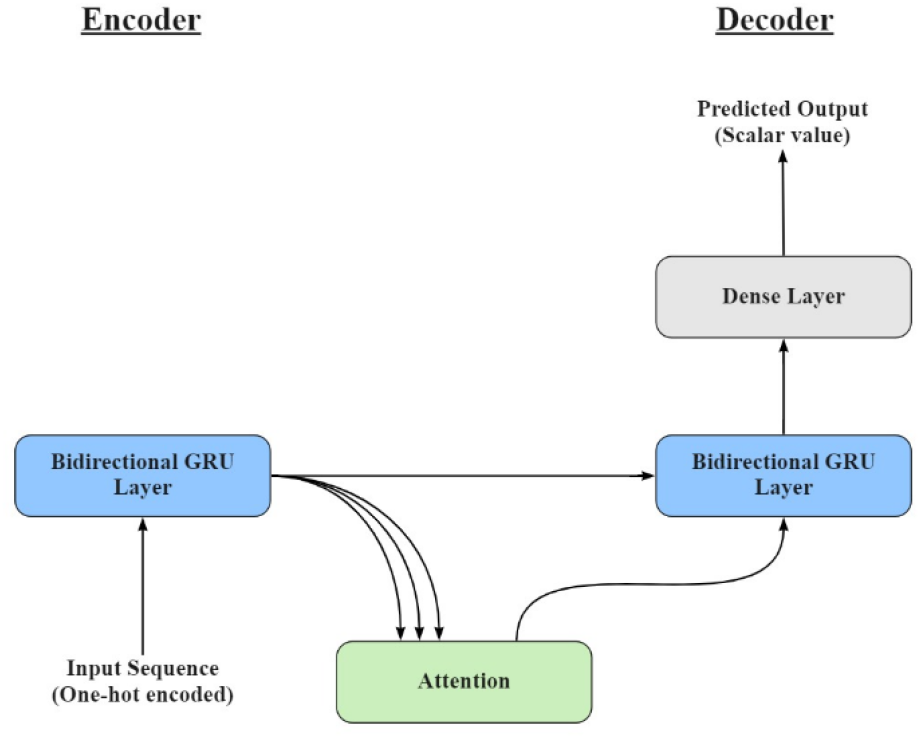
Schematic overview of the neural network architecture, consisting of an autoencoder with attention mechanism. The autoencoder consists of an encoder (one bi-directional GRU layer) and a decoder (one bi-directional GRU layer and a dense layer).

### 4.4. Training and testing

The implementation was done in Python (v.3.8.8) with TensorFlow [91] (v. 2.5.0) using the following libraries: Scikit-learn [85] (0.24.1), Statsmodels [92] (v. 0.12.2), Pandas [93] (v.1.2.4), Matplotlib [94] (v.3.3.4), Seaborn [95] (v.0.11.1), SciPy [96] (v.1.6.2), and NumPy [97] (v.1.20.1). The inputs of the final model are one-hot encoded peptides sequences with a maximum length of 40 residues, where shorter peptides were padded. Thus, the input matrix has a dimension of batch size x 40 x 21, were the 20 AAs and one padding character are included. For the proof-of-concept models, one-hot encoded inputs were also applied. Two formats of sequence were generated with a maximum length of 8 and 40 and input dimensions of batch size x 8 x 10 and batch size x 40 x 21 respectively. Padding was applied to shorter sequences for proof-of-concept models in a same way.

Initially, the proof-of-concept models were trained and optimized to determine their efficacy. Then, the relevant physicochemical properties were determined, by training and optimizing the models using the whole filtered data. The data was randomly split into three sets. A training dataset, which consisted of 90% of the data from which 20% of this data was taken to use as validation data to control overfitting. The remaining 10% of the data was used as test dataset. The test dataset was used to evaluate the generalization capacity, to give an unbiased evaluation of the models and to obtain the results. The three datasets were randomly selected.

To determine the physicochemical properties that influences the MS response for peptides, the attention weights for the 20 AAs were determined. The assigned attention weights for each sequence element were extracted for each intensity prediction. Then those weights were averaged for each AA, first within the same sequence (in case there are repeated AAs within the peptide sequence) and subsequently across all the sequences. The relevancy of the physicochemical properties was determined by computing the Pearson correlation coefficient (PCC) between the average attention weights of AAs and each of the AAindex1 indices.

[30]. An AAindex1 index was considered significant if PCC ≤-0.7 or PCC ≥ 0.7. For the proof-of-concept models the average attention weights were correlated with the fixed contribution assigned to each element of the corresponding sequence format. Once the relevant physicochemical properties were identified, the data was more thoroughly filtered to improve the model’s performance (regarding the MS intensity prediction).

The Mean Squared Error (MSE) was used as the Loss function [98,99]. The accuracy measurement during the models’ training was done using the Mean Absolute Error (MAE), while the models’ performance was expressed with the Mean Absolute Percentage Error (MAPE) [100,101]. Adam [102] was the optimizer chosen, using its default settings. The models were trained on NVIDIA Quadro T2000 GPU for 5 to 30 epochs.

## Data availability

The data used in this project can be found in the PRIDE repository with the identifiers PXD004732, PXD010595, and PXD021013 including the mass spectrometric raw data and the search data.

## Author contribution

**Naim Abdul-Khalek:** Conceptualization, Methodology, Software, Validation, Formal analysis, Investigation, Data Curation, Writing - Original Draft, Writing - Review & Editing, Visualization. **Reinhard Wimmer:** Conceptualization, Resources, Writing - Review & Editing, Supervision. **Michael Toft Overgaard:** Conceptualization, Resources, Writing - Review & Editing, Supervision, Funding acquisition. **Simon Gregersen Echers:** Conceptualization, Methodology, Resources, Validation, Investigation, Writing - Original Draft, Writing - Review & Editing, Supervision, Project administration, Funding acquisition.

## REFERENCES

[1] Awad H, Khamis MM, El-Aneed A. Mass Spectrometry, Review of the Basics: Ionization. Http://DxDoiOrg/101080/057049282014954046 2014;50:158–75. https://doi.org/10.1080/05704928.2014.954046.

[2] Herrero M, Simõ C, García-Cañas V, Ibáñez E, Cifuentes A. Foodomics: MS-based strategies in modern food science and nutrition. Mass Spectrom Rev 2012;31:49–69. https://doi.org/10.1002/MAS.20335.

[3] Davison J, O’Gorman A, Brennan L, Cotter DR. A systematic review of metabolite biomarkers of schizophrenia. Schizophr Res 2018;195:32–50. https://doi.org/10.1016/J.SCHRES.2017.09.021.

[4] Hofstadler SA, Sannes-Lowery KA. Applications of ESI-MS in drug discovery: interrogation of noncovalent complexes. Nature Reviews Drug Discovery 2006 5:7 2006;5:585–95. https://doi.org/10.1038/nrd2083.

[5] García-Moreno PJ, Gregersen S, Nedamani ER, Olsen TH, Marcatili P, Overgaard MT, et al. Identification of emulsifier potato peptides by bioinformatics: application to omega-3 delivery emulsions and release from potato industry side streams. Scientific Reports 2020 10:1 2020;10:1–22. https://doi.org/10.1038/s41598-019-57229-6.

[6] Gregersen S, Kongsted ASH, Nielsen RB, Hansen SS, Lau FA, Rasmussen JB, et al. Enzymatic extraction improves intracellular protein recovery from the industrial carrageenan seaweed Eucheuma denticulatum revealed by quantitative, subcellular protein profiling: A high potential source of functional food ingredients. Food Chem X 2021;12:100137. https://doi.org/10.1016/J.FOCHX.2021.100137.

[7] El-Aneed A, Cohen A, Banoub J. Mass Spectrometry, Review of the Basics: Electrospray, MALDI, and Commonly Used Mass Analyzers. Appl Spectrosc Rev 2009;44:210–30. https://doi.org/10.1080/05704920902717872.

[8] Wilm M. Principles of Electrospray Ionization. Molecular & Cellular Proteomics 2011;10:M111.009407. https://doi.org/10.1074/MCP.M111.009407.

[9] Liuni P, Wilson DJ. Understanding and optimizing electrospray ionization techniques for proteomic analysis. Http://DxDoiOrg/101586/Epr10111 2014;8:197–209. https://doi.org/10.1586/EPR.10.111.

[10] Cañas Montalvo B, López-Ferrer D, Ramos-Fernández A, Camafeita E, Calvo E. Mass spectrometry technologies for proteomics. Brief Funct Genomics 2006;4:295–320. https://doi.org/10.1093/BFGP/ELI002.

[11] Schwanhüusser B, Busse D, Li N, Dittmar G, Schuchhardt J, Wolf J, et al. Global quantification of mammalian gene expression control. Nature 2011;473:337–42. https://doi.org/10.1038/nature10098.

[12] Cox J, Hein MY, Luber CA, Paron I, Nagaraj N, Mann M. Accurate proteome-wide label-free quantification by delayed normalization and maximal peptide ratio extraction, termed MaxLFQ. Molecular and Cellular Proteomics 2014;13:2513–26. https://doi.org/10.1074/mcp.M113.031591.

[13] Nikolov M, Schmidt C, Urlaub H. Quantitative mass spectrometry-based proteomics: An overview. Methods in Molecular Biology 2012;893:85–100. https://doi.org/10.1007/978-1-61779-885-6_7.

[14] Xie F, Liu T, Qian WJ, Petyuk VA, Smith RD. Liquid Chromatography-Mass Spectrometry-based Quantitative Proteomics *. Journal of Biological Chemistry 2011;286:25443–9. https://doi.org/10.1074/JBC.R110.199703.

[15] Vidova V, Spacil Z. A review on mass spectrometry-based quantitative proteomics: Targeted and data independent acquisition. Anal Chim Acta 2017;964:7–23. https://doi.org/10.1016/J.ACA.2017.01.059.

[16] Nahnsen S, Bielow C, Reinert K, Kohlbacher O. Tools for Label-free Peptide Quantification* □ S 2012. https://doi.org/10.1074/mcp.R112.025163.

[17] He B, Shi J, Wang X, Jiang H, Zhu HJ. Label-free absolute protein quantification with data-independent acquisition. J Proteomics 2019;200:51–9. https://doi.org/10.1016/J.JPROT.2019.03.005.

[18] Wiśniewski JR, Hein MY, Cox J, Mann M. A “proteomic ruler” for protein copy number and concentration estimation without spike-in standards. Molecular and Cellular Proteomics 2014;13:3497–506. https://doi.org/10.1074/mcp.M113.037309.

[19] Jafarpour A, Gregersen S, Gomes RM, Marcatili P, Olsen TH, Jacobsen C, et al. Biofunctionality of Enzymatically Derived Peptides from Codfish (Gadus morhua) Frame: Bulk In Vitro Properties, Quantitative Proteomics, and Bioinformatic Prediction. Mar Drugs 2020;18:599. https://doi.org/10.3390/MD18120599.

[20] Gregersen Echers S, Jafarpour A, Yesiltas B, García-Moreno PJ, Greve-Poulsen M, Hansen DK, et al. Targeted hydrolysis of native potato protein: A novel workflow for obtaining hydrolysates with improved interfacial properties. Food Hydrocoll 2023;137:108299. https://doi.org/10.1016/J.FOODHYD.2022.108299.

[21] Wen B, Zeng W-F, Liao Y, Shi Z, Savage SR, Jiang W, et al. Deep Learning in Proteomics. Proteomics 2020;20:1900335. https://doi.org/10.1002/PMIC.201900335.

[22] Chollet F. Deep Learning with Python. 1st ed. USA: Manning Publications Co.; 2017.

[23] Alquraishi M. AlphaFold at CASP13. Bioinformatics 2019;35:4862–5. https://doi.org/10.1093/BIOINFORMATICS/BTZ422.

[24] Sun T, Zhou B, Lai L, Pei J. Sequence-based prediction of protein protein interaction using a deep-learning algorithm. BMC Bioinformatics 2017;18:1–8. https://doi.org/10.1186/S12859-017-1700-2.

[25] Kulmanov M, Hoehndorf R. DeepGOPlus: improved protein function prediction from sequence. Bioinformatics 2020;36:422–9. https://doi.org/10.1093/BIOINFORMATICS/BTZ595.

[26] Meyer JG. Deep learning neural network tools for proteomics. Cell Reports Methods 2021;1:100003. https://doi.org/10.1016/J.CRMETH.2021.100003.

[27] Sonsare PM, Gunavathi C. Investigation of machine learning techniques on proteomics: A comprehensive survey. Prog Biophys Mol Biol 2019;149:54–69. https://doi.org/10.1016/J.PBIOMOLBIO.2019.09.004.

[28] Xu CM, Zhang JY, Liu H, Sun HC, Zhu YP, Xie HW. Advance of Peptide Detectability Prediction on Mass Spectrometry Platform in Proteomics. Chinese Journal of Analytical Chemistry 2010;38:286–92. https://doi.org/10.1016/S1872-2040(09)60023-2.

[29] Gao Z, Chang C, Yang J, Zhu Y, Fu Y. AP3: An Advanced Proteotypic Peptide Predictor for Targeted Proteomics by Incorporating Peptide Digestibility. Anal Chem 2019;91:8705–11. https://doi.org/10.1021/ACS.ANALCHEM.9B02520.

[30] Riley RM, Miko SES, Morin RD, Morin GB, Negri GL. PeptideRanger: An R Package to Optimize Synthetic Peptide Selection for Mass Spectrometry Applications. J Proteome Res 2023;22:526–31. https://doi.org/10.1021/ACS.JPROTEOME.2C00538.

[31] Pauletti BA, Granato DC, M. Carnielli C, Câmara GA, Normando AGC, Telles GP, et al. Typic: A Practical and Robust Tool to Rank Proteotypic Peptides for Targeted Proteomics. J Proteome Res 2023;22:539–45. https://doi.org/10.1021/ACS.JPROTEOME.2C00585.

[32] Rusilowicz M, Newman DW, Creamer DR, Johnson J, Adair K, Harman VM, et al. AlacatDesigner–Computatìonal Design of Peptide Concatamers for Protein Quantitation. J Proteome Res 2023;22:594–604. https://doi.org/10.1021/ACS.JPROTEOME.2C00608.

[33] Mallick P, Schirle M, Chen SS, Flory MR, Lee H, Martin D, et al. Computational prediction of proteotypic peptides for quantitative proteomics. Nat Biotechnol 2006;25:125–31. https://doi.org/10.1038/nbt1275.

[34] Demeure K, Duriez E, Domon B, Niclou SP. Peptide manager: A peptide selection tool for targeted proteomic studies involving mixed samples from different species. Front Genet 2014;5:305. https://doi.org/10.3389/FGENE.2014.00305.

[35] Chen Q, Jiang Y, Ren Y, Ying M, Lu B. Peptide Selection for Accurate Targeted Protein Quantification via a Dimethylation High-Resolution Mass Spectrum Strategy with a Peptide Release Kinetic Model. ACS Omega 2020;5:3809–19. https://doi.org/10.1021/ACSOMEGA.9B02002.

[36] Vaudel M, Burkhart JM, Zahedi RP, Oveland E, Berven FS, Sickmann A, et al. PeptideShaker enables reanalysis of MS-derived proteomics data sets. Nat Biotechnol 2015;33:22–4. https://doi.org/10.1038/nbt.3109.

[37] Rehfeldt TG, Krawczyk K, Bøgebjerg M, Schwammle V, Rottger R. MS2AI: automated repurposing of public peptide LC-MS data for machine learning applications. Bioinformatics 2022;38:875–7. https://doi.org/10.1093/BIOINFORMATICS/BTAB701.

[38] Zolg DP, Wilhelm M, Schnatbaum K, Zerweck J, Knaute T, Delanghe B, et al. Building ProteomeTools based on a complete synthetic human proteome. Nat Methods 2017;14:259–62. https://doi.org/10.1038/NMETH.4153.

[39] Gessulat S, Schmidt T, Zolg DP, Samaras P, Schnatbaum K, Zerweck J, et al. Prosit: proteome-wide prediction of peptide tandem mass spectra by deep learning. Nat Methods 2019;16:509–18. https://doi.org/10.1038/S41592-019-0426-7.

[40] Wilhelm M, Zolg DP, Graber M, Gessulat S, Schmidt T, Schnatbaum K, et al. Deep learning boosts sensitivity of mass spectrometry-based immunopeptidomics. Nature Communications 2021 12:1 2021;12:1–12. https://doi.org/10.1038/s41467-021-23713-9.

[41] Kawashima S, Ogata H, Kanehisa M. AAindex: Amino Acid Index Database. Nucleic Acids Res 1999;27:368–9. https://doi.org/10.1093/NAR/27.1.368.

[42] Anderle M, Roy S, Lin H, Becker C, Joho K. Quantifying reproducibility for differential proteomics: noise analysis for protein liquid chromatography-mass spectrometry of human serum. Bioinformatics 2004;20:3575–82. https://doi.org/10.1093/BIOINFORMATICS/BTH446.

[43] Liu K, Li S, Wang L, Ye Y, Tang H. Full-Spectrum Prediction of Peptides Tandem Mass Spectra using Deep Neural Network. Anal Chem 2020;92:4275–83. https://doi.org/10.1021/ACS.ANALCHEM.9B04867.

[44] Zhou C, Bowler LD, Feng J. A machine learning approach to explore the spectra intensity pattern of peptides using tandem mass spectrometry data. BMC Bioinformatics 2008;9:1–17. https://doi.org/10.1186/1471-2105-9-325.

[45] Cox J, Mann M. MaxQuant enables high peptide identification rates, individualized p.p.b.-range mass accuracies and proteome-wide protein quantification. Nature Biotechnology 2008;26:1367–72. https://doi.org/10.1038/nbt.1511.

[46] Cox J, Neuhauser N, Michalski A, Scheltema RA, Olsen J v., Mann M. Andromeda: A peptide search engine integrated into the MaxQuant environment. J Proteome Res 2011;10:1794–805. https://doi.org/10.1021/PR101065J.

[47] Gregersen S, Pertseva M, Marcatili P, Holdt SL, Jacobsen C, García-Moreno PJ, et al. Proteomic characterization of pilot scale hot-water extracts from the industrial carrageenan red seaweed Eucheuma denticulatum. Algal Res 2022;62:102619. https://doi.org/10.1016/J.ALGAL.2021.102619.

[48] Weinkauf R, Schanen P, Yang D, Soukara / S, Schlag EW. Elementary Processes in Peptides: Electron Mobility and Dissociation in Peptide Cations in the Gas Phase. J Phys Chem 1995;99:11255–65.

[49] Marchese R, Grandori R, Carloni P, Raugei S. On the Zwitterionic Nature of Gas-Phase Peptides and Protein Ions. PLoS Comput Biol 2010;6:e1000775. https://doi.org/10.1371/JOURNAL.PCBI.1000775.

[50] Cech NB, Krone JR, Enke CG. Predicting electrospray response from chromatographic retention time. Anal Chem 2001;73:208–13. https://doi.org/10.1021/AC0006019.

[51] Cech NB, Enke CG. Relating electrospray ionization response to nonpolar character of small peptides. Anal Chem 2000;72:2717–23. https://doi.org/10.1021/AC9914869.

[52] Osaka I, Takayama M. Influence of hydrophobicity on positive-and negative-ion yields of peptides in electrospray ionization mass spectrometry. Rapid Commun Mass Spectrom 2014;28:2222–6. https://doi.org/10.1002/RCM.7010.

[53] Vreeke GJC, Lubbers W, Vincken JP, Wierenga PA. A method to identify and quantify the complete peptide composition in protein hydrolysates. Anal Chim Acta 2022;1201:339616. https://doi.org/10.1016/J.ACA.2022.339616.

[54] Eyers CE, Lawless C, Wedge DC, Lau KW, Gaskell SJ, Hubbard SJ. CONSeQuence: prediction of reference peptides for absolute quantitative proteomics using consensus machine learning approaches. Mol Cell Proteomics 2011;10. https://doi.org/10.1074/MCP.M110.003384.

[55] Zimmer D, Schneider K, Sommer F, Schroda M, Mühlhaus T. Artificial Intelligence Understands Peptide Observability and Assists With Absolute Protein Quantification. Front Plant Sci 2018;9. https://doi.org/10.3389/FPLS.2018.01559.

[56] Muntel J, Boswell SA, Tang S, Ahmed S, Wapinski I, Foley G, et al. Abundance-based classifier for the prediction of mass spectrometric peptide detectability upon enrichment (PPA). Mol Cell Proteomics 2015;14:430–40. https://doi.org/10.1074/MCP.M114.044321.

[57] Qeli E, Omasits U, Goetze S, Stekhoven DJ, Frey JE, Basler K, et al. Improved prediction of peptide detectability for targeted proteomics using a rank-based algorithm and organism-specific data. J Proteomics 2014;108:269–83. https://doi.org/10.1016/J.JPROT.2014.05.011.

[58] Jarnuczak AF, Lee DCH, Lawless C, Holman SW, Eyers CE, Hubbard SJ. Analysis of Intrinsic Peptide Detectability via Integrated Label-Free and SRM-Based Absolute Quantitative Proteomics. J Proteome Res 2016;15:2945–59. https://doi.org/10.1021/ACS.JPROTEOME.6B00048.

[59] Abaye DA, Pullen FS, Nielsen B v. Peptide polarity and the position of arginine as sources of selectivity during positive electrospray ionisation mass spectrometry. Rapid Commun Mass Spectrom 2011;25:3597–608. https://doi.org/10.1002/RCM.5270.

[60] Gautier V, Boumeester AJ, Lössl P, Heck AJR. Lysine conjugation properties in human IgGs studied by integrating high-resolution native mass spectrometry and bottom-up proteomics. Proteomics 2015;15:2756–65. https://doi.org/10.1002/PMIC.201400462.

[61] Searle BC, Egertson JD, Bollinger JG, Stergachis AB, MacCoss MJ. Using Data Independent Acquisition (DIA) to Model High-responding Peptides for Targeted Proteomics Experiments. Molecular & Cellular Proteomics 2015;14:2331–40. https://doi.org/10.1074/MCP.M115.051300.

[62] Mallick P, Schirle M, Chen SS, Flory MR, Lee H, Martin D, et al. Computational prediction of proteotypic peptides for quantitative proteomics. Nat Biotechnol 2006;25:125–31. https://doi.org/10.1038/nbt1275.

[63] Eisenberg D, Weiss RM, Terwilliger TC. The hydrophobic moment detects periodicity in protein hydrophobicity. Proc Natl Acad Sci U S A 1984;81:140–4. https://doi.org/10.1073/PNAS.81.1.140.

[64] Doig AJ, Sternberg MJE. Side-chain conformational entropy in protein folding. Protein Sci 1995;4:2247–51. https://doi.org/10.1002/PRO.5560041101.

[65] Eisenberg D, Mclachlan AD. Solvation energy in protein folding and binding. Nature 1986 319:6050 1986;319:199–203. https://doi.org/10.1038/319199a0.

[66] Jacobs RE, White SH. The nature of the hydrophobic binding of small peptides at the bilayer interface: Implications for the insertion of transbilayer helices. Biochemistry 1989;28:3421–37. https://doi.org/10.1021/BI00434A042.

[67] Zhou H, Zhou Y. Quantifying the effect of burial of amino acid residues on protein stability. Proteins: Structure, Function, and Bioinformatics 2004;54:315–22. https://doi.org/10.1002/PROT.10584.

[68] Novák P, Havlíček V. Protein Extraction and Precipitation. Proteomic Profiling and Analytical Chemistry: The Crossroads: Second Edition 2016:51–62. https://doi.org/10.1016/B978-0-444-63688-1.00004-5.

[69] Finkelstein A V., Badretdinov AY, Ptitsyn OB. Physical reasons for secondary structure stability: alpha-helices in short peptides. Proteins 1991;10:287–99. https://doi.org/10.1002/PROT.340100403.

[70] Marcoux J, Robinson C V. Twenty Years of Gas Phase Structural Biology. Structure 2013;21:1541–50. https://doi.org/10.1016/J.STR.2013.08.002.

[71] Loo JA. Studying noncovalent protein complexes by electrospray ionization mass spectrometry. Mass Spectrom Rev 1998;16:1–23. https://doi.org/10.1002/(SICI)1098-2787(1997)16:1<1::AID-MAS1>3.0.CO;2-L.

[72] Chin W, Compagnon I, Dognon JP, Canuel C, Piuzzi F, Dimicoli I, et al. Spectroscopic evidence for gas-phase formation of successive β-turns in a three-residue peptide chain. J Am Chem Soc 2005;127:1388–9. https://doi.org/10.1021/JA042860B.

[73] Wenschuh H, Halada P, Lamer S, Jungblut P, Krause E. The Ease of Peptide Detection by Matrix-assisted Laser Desorption/Ionization Mass Spectrometry: the Effect of Secondary Structure on Signal Intensity. Rapid Communications in Mass Spectrometry 1998;12:115–9. https://doi.org/10.1002/(SICI)1097-0231(19980214)12:3<115::AID-RCM124>3.0.CO;2-5.

[74] Tsaprailis G, Nair H, Somogyi Á, Wysocki VH, Zhong W, Futrell JH, et al. Influence of secondary structure on the fragmentation of protonated peptides. J Am Chem Soc 1999;121:5142–54. https://doi.org/10.1021/JA982980H.

[75] Ruotolo BT, Verbeck GF, Thomson LM, Gillig KJ, Russell DH. Observation of conserved solution-phase secondary structure in gas-phase tryptic peptides. J Am Chem Soc 2002;124:4214–5. https://doi.org/10.1021/JA0178113.

[76] Ruotolo BT, Russell DH. Gas-Phase Conformations of Proteolytically Derived Protein Fragments: Influence of Solvent on Peptide Conformation. Journal of Physical Chemistry B 2004;108:15321–31. https://doi.org/10.1021/JP0490296.

[77] Bonfiglio R, King RC, Olah T v, Merkle K. The Effects of Sample Preparation Methods on the Variability of the Electrospray Ionization Response for Model Drug Compounds. Rapid Communications in Mass Spectrometry 1999;13:1175–85. https://doi.org/10.1002/(SICI)1097-0231(19990630)13:12<1175::AID-RCM639>3.0.CO;2-0.

[78] Šedo O, Sedláček I, Zdráhal Z. Sample preparation methods for MALDI-MS profiling of bacteria. Mass Spectrom Rev 2011;30:417–34. https://doi.org/10.1002/MAS.20287.

[79] Nilsson T, Mann M, Aebersold R, Yates JR, Bairoch A, Bergeron JJM. Mass spectrometry in high-throughput proteomics: ready for the big time. Nat Methods 2010;7:681–5. https://doi.org/10.1038/nmeth0910-681.

[80] Tabb DL, Vega-Montoto L, Rudnick PA, Variyath AM, Ham AJL, Bunk DM, et al. Repeatability and reproducibility in proteomic identifications by liquid chromatography-tandem mass spectrometry. J Proteome Res 2009;9:761–76. https://doi.org/10.1021/PR9006365.

[81] Haag AM. Mass Analyzers and Mass Spectrometers BT - Modern Proteomics – Sample Preparation, Analysis and Practical Applications. In: Mirzaei H, Carrasco M, editors., Cham: Springer International Publishing; 2016, p. 157–69. https://doi.org/10.1007/978-3-319-41448-5_7.

[82] Nordström A, Want E, Northen T, Lehtiö J, Siuzdak G. Multiple ionization mass spectrometry strategy used to reveal the complexity of metabolomics. Anal Chem 2008;80:421–9. https://doi.org/10.1021/AC701982E.

[83] Bell AW, Deutsch EW, Au CE, Kearney RE, Beavis R, Sechi S, et al. A HUPO test sample study reveals common problems in mass spectrometry–based proteomics. Nature Methods 2009 6:6 2009;6:423–30. https://doi.org/10.1038/nmeth.1333.

[84] Boutilier K, Ross M, Podtelejnikov A v., Orsi C, Taylor R, Taylor P, et al. Comparison of different search engines using validated MS/MS test datasets. Anal Chim Acta 2005;534:11–20. https://doi.org/10.1016/J.ACA.2004.04.047.

[85] Pedregosa F, Varoquaux G, Gramfort A, Michel V, Thirion B, Grisel O, et al. Scikit-learn: Machine Learning in Python. Journal of Machine Learning Research 2012;12:2825–30. https://doi.org/10.48550/arxiv.1201.0490.

[86] Silva ASC, Bouwmeester R, Martens L, Degroeve S. Accurate peptide fragmentation predictions allow data driven approaches to replace and improve upon proteomics search engine scoring functions. Bioinformatics 2019;35:5243–8. https://doi.org/10.1093/BIOINFORMATICS/BTZ383.

[87] Bowden P, Thavarajah T, Zhu P, McDonell M, Thiele H, Marshall JG. Quantitative statistical analysis of standard and human blood proteins from liquid chromatography, electrospray ionization, and tandem mass spectrometry. J Proteome Res 2012;11:2032–47. https://doi.org/10.1021/PR2000013.

[88] Ryu S, Goodlett DR, Noble WS, Minin VN. A statistical approach to peptide identification from clustered tandem mass spectrometry data. 2012 IEEE International Conference on Bioinformatics and Biomedicine Workshops 2012:648–53. https://doi.org/10.1109/BIBMW.2012.6470214.

[89] Chung J, Gulcehre C, Cho K, Bengio Y. Empirical Evaluation of Gated Recurrent Neural Networks on Sequence Modeling 2014. https://doi.org/10.48550/arxiv.1412.3555.

[90] Bahdanau D, Cho KH, Bengio Y. Neural Machine Translation by Jointly Learning to Align and Translate. 3rd International Conference on Learning Representations, ICLR 2015 - Conference Track Proceedings 2014. https://doi.org/10.48550/arxiv.1409.0473.

[91] Abadi M, Barham P, Chen J, Chen Z, Davis A, Dean J, et al. TensorFlow: A system for large-scale machine learning. Proceedings of the 12th USENIX Symposium on Operating Systems Design and Implementation, OSDI 2016 2016:265–83. https://doi.org/10.48550/arxiv.1605.08695.

[92] Seabold S, Perktold J. Statsmodels: Econometric and Statistical Modeling with Python. Proceedings of the 9th Python in Science Conference 2010:92–6. https://doi.org/10.25080/MAJORA-92BF1922-011.

[93] McKinney W. Data Structures for Statistical Computing in Python. Proceedings of the 9th Python in Science Conference 2010:56–61. https://doi.org/10.25080/MAJORA-92BF1922-00A.

[94] Hunter JD. Matplotlib: A 2D graphics environment. Comput Sci Eng 2007;9:90–5. https://doi.org/10.1109/MCSE.2007.55.

[95] Waskom ML. seaborn: statistical data visualization. J Open Source Softw 2021;6:3021. https://doi.org/10.21105/JOSS.03021.

[96] Virtanen P, Gommers R, Oliphant TE, Haberland M, Reddy T, Cournapeau D, et al. SciPy 1.0: Fundamental algorithms for scientific computing in Python. Nat Methods 2020;17:261–72. https://doi.org/10.1038/s41592-019-0686-2.

[97] Harris CR, Millman KJ, van der Walt SJ, Gommers R, Virtanen P, Cournapeau D, et al. Array Programming with NumPy. Nature 2020;585:357–62. https://doi.org/10.1038/s41586-020-2649-2.

[98] Shimobaba T, Kakue T, Ito T. Convolutional Neural Network-Based Regression for Depth Prediction in Digital Holography. 2018 IEEE 27th International Symposium on Industrial Electronics 2018:1323–6. https://doi.org/10.1109/ISIE.2018.8433651.

[99] Park A, Joo M, Kim K, Son WJ, Lim GT, Lee J, et al. A comprehensive evaluation of regression-based drug responsiveness prediction models, using cell viability inhibitory concentrations (IC50 values). Bioinformatics 2022;38:2810–7. https://doi.org/10.1093/BIOINFORMATICS/BTAC177.

[100] Nguyen M, Jankovic I, Kalesinskas L, Baiocchi M, Chen JH. Machine learning for initial insulin estimation in hospitalized patients. Journal of the American Medical Informatics Association 2021;28:2212–9. https://doi.org/10.1093/JAMIA/OCAB099.

[101] Ren Y, Li X, Xu H. A Deep Learning Model to Extract Ship Size from Sentinel-1 SAR Images. IEEE Transactions on Geoscience and Remote Sensing 2022;60:1–14. https://doi.org/10.1109/TGRS.2021.3063216.

[102] Kingma DP, Ba JL. Adam: A Method for Stochastic Optimization. 3rd International Conference on Learning Representations 2014. https://doi.org/10.48550/arxiv.1412.6980.

