## Supplementary information for "Insight on physicochemical properties governing peptide MS1 response in HPLC-ESI-MS/MS proteomics: A deep learning approach"

### for

**Table S1**. Example of element contribution values for proof-of-concept models' data design. For instance, the sequence ACCBAADAEA will have an output value of 100 (1 + 10 + 10 + 5 + 1 + 1 + 20 + 1 + 50 + 1 = 100).

| Element | Value |
|---|---|
| **A** | 1 |
| **B** | 5 |
| **C** | 10 |
| **D** | 20 |
| **E** | 50 |

**Table S2**. Element contribution values and obtained attention weights for proof-of-concept models 1 and 2.

| | Proof-of-concept Model 1 | | Proof-of-concept Model 2 | |
|---|---|---|---|---|
| **Element** | **Value** | **Attention Weight** | **Value** | **Attention Weight** |
| **1** | 10 | 0,123905 | 1 | 0,081468 |
| **2** | 20 | 0,124513 | 10 | 0,112548 |
| **3** | 30 | 0,124988 | 3 | 0,08754 |
| **4** | 40 | 0,125741 | 40 | 0,196395 |
| **5** | 50 | 0,126275 | 5 | 0,093186 |
| **6** | 60 | 0,126778 | 90 | 0,282256 |
| **7** | 70 | 0,12751 | 7 | 0,101281 |
| **8** | 80 | 0,128112 | 160 | 0,350464 |
| **9** | 90 | 0,128624 | 9 | 0,110186 |

**Table S3**. Proof-of-concept models 1 and 2 results showing correlation of attention weights and contribution values (PCC and associated p-value), performance metric (MAPE), and hyperparameter specifications.

| | PCC[a] | p-value | MAPE[b] (%) | Batch Size | Units | Epochs | Output range of values |
|---|---|---|---|---|---|---|---|
| **Proof-of-concept Model 1** | 0.999 | <3E-11 | 0.79 | 8 | 128 | 5 | 1-12 |
| **Proof-of-concept Model 2** | 0.980 | <4E-06 | 0.56 | 8 | 64 | 20 | 1-12 |

[a] Pearson correlation coefficient

[b] Mean absolute percentage error

**Table S4**. Element contribution values and obtained attention weights for proof-of-concept models 3 and 4.

| | **Proof-of-concept Model 3** | | **Proof-of-concept Model 4** | |
|---|---|---|---|---|
| **Element** | **Value** | **Attention Weight** | **Value** | **Attention Weight** |
| **1** | $1,0x10^{7}$ | 0,02216 | 0 | 0,004297 |
| **2** | $2,0x10^{7}$ | 0,024845 | $6,0x10^{6}$ | 0,009937 |
| **3** | $3,0x10^{7}$ | 0,0262 | 3 | 0,003927 |
| **4** | $4,0x10^{7}$ | 0,029228 | $1,2x10^{7}$ | 0,014019 |
| **5** | $5,0x10^{7}$ | 0,031261 | 10 | 0,004406 |
| **6** | $6,0x10^{7}$ | 0,033604 | $1,8x10^{7}$ | 0,017216 |
| **7** | $7,0x10^{7}$ | 0,03581 | 21 | 0,004617 |
| **8** | $8,0x10^{7}$ | 0,039363 | $2,4x10^{7}$ | 0,01984 |
| **9** | $9,0x10^{7}$ | 0,041557 | 36 | 0,004456 |
| **10** | $1,0x\ 10^{8}$ | 0,043841 | $3,0x10^{7}$ | 0,022459 |
| **11** | $1,1x10^{8}$ | 0,045409 | $1,1x10^{6}$ | 0,005686 |
| **12** | $1,2x10^{8}$ | 0,048081 | $7,2x10^{8}$ | 0,124603 |
| **13** | $1,3x10^{8}$ | 0,051647 | $1,3x10^{6}$ | 0,005725 |
| **14** | $1,4x10^{8}$ | 0,052853 | $8,4x10^{8}$ | 0,132533 |
| **15** | $1,5x10^{8}$ | 0,056963 | $1,5x10^{6}$ | 0,005524 |
| **16** | $1,6x10^{8}$ | 0,060133 | $1,28x10^{9}$ | 0,164589 |
| **17** | $1,7x10^{8}$ | 0,06256 | $1,7x10^{6}$ | 0,006098 |
| **18** | $1,8x10^{8}$ | 0,064004 | $1,44x10^{9}$ | 0,174379 |
| **19** | $1,9x10^{8}$ | 0,06645 | $1,9x10^{6}$ | 0,006145 |
| **20** | $2,0x10^{8}$ | 0,067933 | $2,0x10^{9}$ | 0,203727 |

**Table S5**. Proof-of-concept models 3 and 4 results showing correlation of attention weights and contribution values (PCC and associated p-value), performance metric (MAPE), and hyperparameter specifications

| | **PCC**[a] | **p-value** | **MAPE**[b] **(%)** | **Batch Size** | **Units** | **Epochs** | **Output range of values** |
|---|---|---|---|---|---|---|---|
| **Proof-of-concept Model 3** | 0.999 | <3E-25 | 0.60 | 32 | 128 | 24 | 1-12 |
| **Proof-of-concept Model 4** | 0.981 | <4E-14 | 3.15 | 32 | 256 | 30 | 1-12 |

[a] Pearson correlation coefficient

[b] Mean absolute percentage error

**Table S6**. Attention weights of individual amino acids (in descending order) for representative model 1 (Bulky hydrophobic and aromatic amino acids).

| Amino Acid | Attention Weight |
|---|---|
| W | 0,158803 |
| L | 0,140030 |
| F | 0,126985 |
| I | 0,102967 |
| Y | 0,096798 |
| V | 0,083330 |
| P | 0,068007 |
| M | 0,059155 |
| C | 0,035384 |
| R | 0,032246 |
| A | 0,025734 |
| D | 0,021801 |
| N | 0,021531 |
| E | 0,021508 |
| K | 0,021058 |
| H | 0,020838 |
| T | 0,019911 |
| S | 0,019147 |
| Q | 0,018424 |
| G | 0,017545 |

**Table S7**. Relevant properties identified by computing the correlation between attention weights of representative model 1 (Bulky hydrophobic and aromatic amino acids) and the AAindex1.

| **Accession number** | **Data description** | **Correlation Score** | **p-value** |
|---|---|---|---|
| MEEJ810102 | Retention coefficient in NaH2PO4 (Meek-Rossetti, 1981) | 0,94 | 4,1E-10 |
| MEEJ810101 | Retention coefficient in NaClO4 (Meek-Rossetti, 1981) | 0,94 | 8,3E-10 |
| BULH740101 | Transfer free energy to surface (Bull-Breese, 1974) | -0,93 | 1,7E-09 |
| GUOD860101 | Retention coefficient at pH 2 (Guo et al., 1986) | 0,93 | 3,0E-09 |
| PARJ860101 | HPLC parameter (Parker et al., 1986) | -0,93 | 4,1E-09 |
| NOZY710101 | Transfer energy, organic solvent/water (Nozaki-Tanford, 1971) | 0,92 | 8,1E-09 |
| WOLS870101 | Principal property value z1 (Wold et al., 1987) | -0,92 | 9,0E-09 |
| MEEJ800102 | Retention coefficient in HPLC, pH2.1 (Meek, 1980) | 0,92 | 1,5E-08 |
| ZHOH040101 | The stability scale from the knowledge-based atom-atom potential (Zhou-Zhou, | 0,92 | 1,5E-08 |
| VENT840101 | Bitterness (Venanzi, 1984) | 0,91 | 3,8E-08 |
| TAKK010101 | Side-chain contribution to protein stability (kJ/mol) (Takano-Yutani, 2001) | 0,89 | 1,1E-07 |
| ZHOH040102 | The relative stability scale extracted from mutation experiments (Zhou-Zhou, | 0,89 | 1,7E-07 |
| PLIV810101 | Partition coefficient (Pliska et al., 1981) | 0,89 | 1,9E-07 |

| | | | |
|---|---|---|---|
| GOLD730101 | Hydrophobicity factor (Goldsack-Chalifoux, 1973) | 0,88 | 2,2E-07 |
| SIMZ760101 | Transfer free energy (Simon, 1976), Cited by Charton-Charton (1982) | 0,88 | 2,8E-07 |
| ARGP820101 | Hydrophobicity index (Argos et al., 1982) | 0,88 | 3,2E-07 |
| JOND750101 | Hydrophobicity (Jones, 1975) | 0,88 | 3,3E-07 |
| ROSM880104 | Hydropathies of amino acid side chains, neutral form (Roseman, 1988) | 0,88 | 1,7E-06 |
| BROC820101 | Retention coefficient in TFA (Browne et al., 1982) | 0,87 | 5,4E-07 |
| CIDH920102 | Normalized hydrophobicity scales for beta-proteins (Cid et al., 1992) | 0,87 | 8,4E-07 |
| ZIMJ680105 | RF rank (Zimmerman et al., 1968) | 0,86 | 1,1E-06 |
| CIDH920105 | Normalized average hydrophobicity scales (Cid et al., 1992) | 0,85 | 1,8E-06 |
| BASU050102 | Interactivity scale obtained by maximizing the mean of correlation | 0,85 | 2,6E-06 |
| MIYS990101 | Relative partition energies derived by the Bethe approximation | -0,84 | 2,9E-06 |
| ZHOH040103 | Buriability (Zhou-Zhou, 2004) | 0,84 | 3,2E-06 |
| MIYS990102 | Optimized relative partition energies - method A (Miyazawa-Jernigan, 1999) | -0,84 | 3,3E-06 |
| WILM950101 | Hydrophobicity coefficient in RP-HPLC, C18 with 0.1%TFA/MeCN/H2O (Wilce et | 0,84 | 3,4E-06 |
| RADA880102 | Transfer free energy from oct to wat (Radzicka-Wolfenden, 1988) | 0,84 | 3,8E-06 |
| FAUJ830101 | Hydrophobic parameter pi (Fauchere-Pliska, 1983) | 0,83 | 4,9E-06 |
| LEVM760106 | van der Waals parameter R0 (Levitt, 1976) | 0,83 | 5,3E-06 |
| WEBA780101 | RF value in high salt chromatography (Weber-Lacey, 1978) | -0,83 | 7,2E-06 |
| BROC820102 | Retention coefficient in HFBA (Browne et al., 1982) | 0,82 | 8,8E-06 |
| CORJ870102 | SWEIG index (Cornette et al., 1987) | 0,81 | 1,2E-05 |
| SWER830101 | Optimal matching hydrophobicity (Sweet-Eisenberg, 1983) | 0,81 | 1,3E-05 |
| BLAS910101 | Scaled side chain hydrophobicity values (Black-Mould, 1991) | 0,81 | 1,4E-05 |
| GUYH850103 | Apparent partition energies calculated from Robson-Osguthorpe index (Guy, | -0,81 | 2,5E-05 |
| LEVM760107 | van der Waals parameter epsilon (Levitt, 1976) | 0,81 | 1,6E-05 |
| ROBB790101 | Hydration free energy (Robson-Osguthorpe, 1979) | 0,81 | 1,7E-05 |
| CIDH920104 | Normalized hydrophobicity scales for alpha/beta-proteins (Cid et al., 1992) | 0,81 | 1,8E-05 |
| WILM950102 | Hydrophobicity coefficient in RP-HPLC, C8 with 0.1%TFA/MeCN/H2O (Wilce et al. | 0,81 | 1,8E-05 |
| MIYS850101 | Effective partition energy (Miyazawa-Jernigan, 1985) | 0,80 | 2,3E-05 |
| ZIMJ680102 | Bulkiness (Zimmerman et al., 1968) | 0,80 | 2,4E-05 |
| CIDH920103 | Normalized hydrophobicity scales for alpha+beta-proteins (Cid et al., 1992) | 0,80 | 2,4E-05 |
| BASU050101 | Interactivity scale obtained from the contact matrix (Bastolla et al., 2005) | 0,80 | 2,5E-05 |
| OOBM770103 | Long range non-bonded energy per atom (Oobatake-Ooi, 1977) | -0,80 | 2,8E-05 |
| EISD860101 | Solvation free energy (Eisenberg-McLachlan, 1986) | 0,79 | 3,3E-05 |
| MEEJ800101 | Retention coefficient in HPLC, pH7.4 (Meek, 1980) | 0,79 | 4,0E-05 |
| GRAR740102 | Polarity (Grantham, 1974) | -0,78 | 5,3E-05 |
| COWR900101 | Hydrophobicity index, 3.0 pH (Cowan-Whittaker, 1990) | 0,78 | 5,7E-05 |
| NISK860101 | 14 A contact number (Nishikawa-Ooi, 1986) | 0,78 | 5,8E-05 |
| ROSG850101 | Mean area buried on transfer (Rose et al., 1985) | 0,77 | 6,3E-05 |
| ROSM880105 | Hydropathies of amino acid side chains, pi-values in pH 7.0 (Roseman, 1988) | 0,77 | 1,1E-04 |
| BASU050103 | Interactivity scale obtained by maximizing the mean of correlation | 0,77 | 7,1E-05 |
| GARJ730101 | Partition coefficient (Garel et al., 1973) | 0,77 | 7,5E-05 |
| PONP800107 | Accessibility reduction ratio (Ponnuswamy et al., 1980) | 0,77 | 7,7E-05 |
| CIDH920101 | Normalized hydrophobicity scales for alpha-proteins (Cid et al., 1992) | 0,77 | 8,4E-05 |

| | | | |
|---|---|---|---|
| KIDA850101 | Hydrophobicity-related index (Kidera et al., 1985) | -0,76 | 9,3E-05 |
| WIMW960101 | Free energies of transfer of AcWl-X-LL peptides from bilayer interface to | 0,76 | 9,6E-05 |
| MIYS990105 | Optimized relative partition energies - method D (Miyazawa-Jernigan, 1999) | -0,76 | 1,0E-04 |
| MEIH800101 | Average reduced distance for C-alpha (Meirovitch et al., 1980) | -0,76 | 1,1E-04 |
| VINM940102 | Normalized flexibility parameters (B-values) for each residue surrounded by | -0,76 | 1,1E-04 |
| MIYS990104 | Optimized relative partition energies - method C (Miyazawa-Jernigan, 1999) | -0,76 | 1,1E-04 |
| BIOV880101 | Information value for accessibility; average fraction 35% (Biou et al., 1988) | 0,75 | 1,2E-04 |
| LEVM760101 | Hydrophobic parameter (Levitt, 1976) | -0,75 | 1,4E-04 |
| BULH740102 | Apparent partial specific volume (Bull-Breese, 1974) | 0,75 | 1,5E-04 |
| OOBM770104 | Average non-bonded energy per residue (Oobatake-Ooi, 1977) | -0,75 | 1,6E-04 |
| BIOV880102 | Information value for accessibility; average fraction 23% (Biou et al., 1988) | 0,74 | 1,9E-04 |
| WERD780101 | Propensity to be buried inside (Wertz-Scheraga, 1978) | 0,73 | 2,3E-04 |
| HOPT810101 | Hydrophilicity value (Hopp-Woods, 1981) | -0,73 | 2,4E-04 |
| LAWE840101 | Transfer free energy, CHP/water (Lawson et al., 1984) | 0,73 | 2,7E-04 |
| NAKH900110 | Normalized composition of membrane proteins (Nakashima et al., 1990) | 0,73 | 2,7E-04 |
| AVBF000109 | Slopes proteins, FDPB VFF neutral (Avbelj, 2000) | 0,73 | 4,3E-04 |
| ROSM880102 | Side chain hydropathy, corrected for solvation (Roseman, 1988) | -0,72 | 3,2E-04 |
| KARP850101 | Flexibility parameter for no rigid neighbors (Karplus-Schulz, 1985) | -0,72 | 3,3E-04 |
| GUYH850102 | Apparent partition energies calculated from Wertz-Scheraga index (Guy, 1985) | -0,72 | 3,3E-04 |
| MIYS990103 | Optimized relative partition energies - method B (Miyazawa-Jernigan, 1999) | -0,72 | 3,3E-04 |
| RACS770101 | Average reduced distance for C-alpha (Rackovsky-Scheraga, 1977) | -0,72 | 3,6E-04 |
| RADA880108 | Mean polarity (Radzicka-Wolfenden, 1988) | 0,71 | 4,2E-04 |
| PUNT030102 | Knowledge-based membrane-propensity scale from 3D_Helix in MPtopo databases | -0,71 | 4,5E-04 |
| RICJ880107 | Relative preference value at N4 (Richardson-Richardson, 1988) | 0,71 | 4,9E-04 |
| PARS000101 | p-Values of mesophilic proteins based on the distributions of B values | -0,71 | 5,0E-04 |
| ROSM880101 | Side chain hydropathy, uncorrected for solvation (Roseman, 1988) | -0,70 | 5,3E-04 |

**Table S8**. Attention weights of individual amino acids (in descending order) for representative model 2 (Positively charged amino acids).

| Amino Acids | Attention Weights |
|---|---|
| R | 0,378370 |
| K | 0,366475 |
| H | 0,172637 |
| P | 0,044171 |
| L | 0,043256 |
| G | 0,042278 |
| S | 0,038368 |
| A | 0,037726 |
| I | 0,037630 |
| N | 0,032570 |
| W | 0,032282 |
| V | 0,031488 |
| F | 0,029724 |
| C | 0,029234 |
| Q | 0,029042 |
| M | 0,026585 |
| D | 0,026067 |
| E | 0,025886 |
| Y | 0,023940 |
| T | 0,023057 |

**Table S9**. Relevant properties identified by computing the correlation between attention weights of representative model 2 (Positively charged amino acids) and the AAindex1.

| Accession number | Data description | Correlation Score | p-value |
|---|---|---|---|
| FAUJ880111 | Positive charge (Fauchere et al., 1988) | 0,93 | 1,6E-09 |
| ZIMJ680104 | Isoelectric point (Zimmerman et al., 1968) | 0,88 | 4,4E-07 |
| EISD860102 | Atom-based hydrophobic moment (Eisenberg-McLachlan, 1986) | 0,84 | 3,5E-06 |
| FINA910103 | Helix termination parameter at posision j-2,j-1,j (Finkelstein et al., 1991) | 0,78 | 4,9E-05 |
| HUTJ700103 | Entropy of formation (Hutchens, 1970) | 0,77 | 7,6E-05 |
| JACR890101 | Weights from the IFH scale (Jacobs-White, 1989) | -0,76 | 1,0E-04 |
| RADA880107 | Energy transfer from out to in(95%buried) (Radzicka-Wolfenden, 1988) | -0,75 | 1,6E-04 |
| KLEP840101 | Net charge (Klein et al., 1984) | 0,74 | 1,8E-04 |
| JANJ780101 | Average accessible surface area (Janin et al., 1978) | 0,72 | 3,5E-04 |

**Table S10**. Performance metrics (expressed as MAPE (%) on real and log-transformed scale) and hyperparameter specifications for the two representative models and the final model (specific/tryptic data subset).

| | **MAPE[a] (%)** | **MAPE[a] (%) Log-Transformed** | **Batch Size** | **Units** | **Epochs** | **Output range of values** |
|---|---|---|---|---|---|---|
| **Representative Model 1** | 165.18 | 12.85 | 128 | 128 | 5 | 1-5 |
| **Representative Model 2** | 231.04 | 16.90 | 512 | 256 | 5 | 1-24 |
| **Final Model** | 85.42 | 10.48 | 256 | 128 | 10 | 1-5 |

[a] Mean absolute percentage error

**Table S11**. Attention weights of individual amino acids (descending weight) for the final model.

| Amino Acid | Attention Weight |
|---|---|
| W | 0,151849 |
| F | 0,119770 |
| L | 0,112485 |
| Y | 0,099900 |
| I | 0,094913 |
| P | 0,088325 |
| V | 0,084304 |
| N | 0,059511 |
| M | 0,056819 |
| T | 0,052511 |
| E | 0,046911 |
| A | 0,046023 |
| R | 0,045235 |
| K | 0,045145 |
| Q | 0,044501 |
| C | 0,043824 |
| S | 0,041541 |
| D | 0,039935 |
| H | 0,039882 |
| G | 0,031428 |

**Table S12**. Relevant properties identified by computing the correlation between attention weights of the final model and AAindex1.

| Accession number | Data description | Correlation Score | p-value |
|---|---|---|---|
| MEEJ800102 | Retention coefficient in HPLC, pH2.1 (Meek, 1980) | 0,91 | 2,6E-08 |
| MEEJ810102 | Retention coefficient in NaH2PO4 (Meek-Rossetti, 1981) | 0,90 | 4,3E-08 |
| NOZY710101 | Transfer energy, organic solvent/water (Nozaki-Tanford, 1971) | 0,90 | 6,2E-08 |
| TAKK010101 | Side-chain contribution to protein stability (kJ/mol) (Takano-Yutani, 2001) | 0,90 | 8,7E-08 |
| GOLD730101 | Hydrophobicity factor (Goldsack-Chalifoux, 1973) | 0,89 | 1,3E-07 |
| MEEJ810101 | Retention coefficient in NaClO4 (Meek-Rossetti, 1981) | 0,88 | 2,3E-07 |
| BULH740101 | Transfer free energy to surface (Bull-Breese, 1974) | -0,88 | 4,4E-07 |
| ZHOH040101 | The stability scale from the knowledge-based atom-atom potential (Zhou-Zhou, | 0,87 | 4,6E-07 |
| ZHOH040102 | The relative stability scale extracted from mutation experiments (Zhou-Zhou, | 0,87 | 5,8E-07 |
| VENT840101 | Bitterness (Venanzi, 1984) | 0,87 | 6,5E-07 |
| WOLS870101 | Principal property value z1 (Wold et al., 1987) | -0,87 | 7,3E-07 |
| SIMZ760101 | Transfer free energy (Simon, 1976), Cited by Charton-Charton (1982) | 0,87 | 7,3E-07 |
| GUOD860101 | Retention coefficient at pH 2 (Guo et al., 1986) | 0,87 | 7,9E-07 |
| ARGP820101 | Hydrophobicity index (Argos et al., 1982) | 0,86 | 9,4E-07 |
| JOND750101 | Hydrophobicity (Jones, 1975) | 0,86 | 9,4E-07 |
| WEBA780101 | RF value in high salt chromatography (Weber-Lacey, 1978) | -0,86 | 1,5E-06 |
| PARJ860101 | HPLC parameter (Parker et al., 1986) | -0,85 | 1,9E-06 |
| LEVM760107 | van der Waals parameter epsilon (Levitt, 1976) | 0,85 | 2,0E-06 |
| ZIMJ680105 | RF rank (Zimmerman et al., 1968) | 0,84 | 3,4E-06 |
| BROC820101 | Retention coefficient in TFA (Browne et al., 1982) | 0,84 | 3,9E-06 |
| GARJ730101 | Partition coefficient (Garel et al., 1973) | 0,82 | 1,0E-05 |
| BROC820102 | Retention coefficient in HFBA (Browne et al., 1982) | 0,82 | 1,1E-05 |
| WILM950101 | Hydrophobicity coefficient in RP-HPLC, C18 with 0.1%TFA/MeCN/H2O (Wilce et | 0,81 | 1,3E-05 |
| ZIMJ680102 | Bulkiness (Zimmerman et al., 1968) | 0,81 | 1,8E-05 |
| PLIV810101 | Partition coefficient (Pliska et al., 1981) | 0,81 | 1,8E-05 |
| RADA880102 | Transfer free energy from oct to wat (Radzicka-Wolfenden, 1988) | 0,80 | 2,2E-05 |
| CIDH920102 | Normalized hydrophobicity scales for beta-proteins (Cid et al., 1992) | 0,80 | 2,5E-05 |
| LEVM760106 | van der Waals parameter R0 (Levitt, 1976) | 0,79 | 2,9E-05 |
| ROSM880104 | Hydropathies of amino acid side chains, neutral form (Roseman, 1988) | 0,79 | 8,8E-05 |
| WILM950102 | Hydrophobicity coefficient in RP-HPLC, C8 with 0.1%TFA/MeCN/H2O (Wilce et al. | 0,79 | 3,5E-05 |
| MEEJ800101 | Retention coefficient in HPLC, pH7.4 (Meek, 1980) | 0,79 | 4,0E-05 |
| AVBF000109 | Slopes proteins, FDPB VFF neutral (Avbelj, 2000) | 0,78 | 9,7E-05 |
| CIDH920105 | Normalized average hydrophobicity scales (Cid et al., 1992) | 0,77 | 6,8E-05 |
| ZHOH040103 | Buriability (Zhou-Zhou, 2004) | 0,77 | 6,9E-05 |
| FAUJ830101 | Hydrophobic parameter pi (Fauchere-Pliska, 1983) | 0,77 | 7,0E-05 |
| OOBM770104 | Average non-bonded energy per residue (Oobatake-Ooi, 1977) | -0,77 | 7,3E-05 |
| BASU050102 | Interactivity scale obtained by maximizing the mean of correlation | 0,76 | 9,6E-05 |
| BLAS910101 | Scaled side chain hydrophobicity values (Black-Mould, 1991) | 0,76 | 1,0E-04 |
| EISD860101 | Solvation free energy (Eisenberg-McLachlan, 1986) | 0,75 | 1,2E-04 |

| | | | |
|---|---|---|---|
| LEVM760101 | Hydrophobic parameter (Levitt, 1976) | -0,74 | 1,7E-04 |
| WIMW960101 | Free energies of transfer of AcWl-X-LL peptides from bilayer interface to | 0,74 | 2,1E-04 |
| ROSG850101 | Mean area buried on transfer (Rose et al., 1985) | 0,73 | 2,3E-04 |
| CORJ870102 | SWEIG index (Cornette et al., 1987) | 0,73 | 2,6E-04 |
| GUYH850103 | Apparent partition energies calculated from Robson-Osguthorpe index (Guy, | -0,73 | 3,9E-04 |
| SWER830101 | Optimal matching hydrophobicity (Sweet-Eisenberg, 1983) | 0,73 | 2,7E-04 |
| MIYS990101 | Relative partition energies derived by the Bethe approximation | -0,73 | 2,8E-04 |
| MIYS990102 | Optimized relative partition energies - method A (Miyazawa-Jernigan, 1999) | -0,73 | 3,0E-04 |
| ROSM880105 | Hydropathies of amino acid side chains, pi-values in pH 7.0 (Roseman, 1988) | 0,72 | 4,9E-04 |
| ROBB790101 | Hydration free energy (Robson-Osguthorpe, 1979) | 0,72 | 3,3E-04 |
| CIDH920104 | Normalized hydrophobicity scales for alpha/beta-proteins (Cid et al., 1992) | 0,72 | 3,5E-04 |
| HOPT810101 | Hydrophilicity value (Hopp-Woods, 1981) | -0,71 | 4,1E-04 |
| OOBM770103 | Long range non-bonded energy per atom (Oobatake-Ooi, 1977) | -0,71 | 4,4E-04 |
| COWR900101 | Hydrophobicity index, 3.0 pH (Cowan-Whittaker, 1990) | 0,71 | 4,7E-04 |
| KIDA850101 | Hydrophobicity-related index (Kidera et al., 1985) | -0,71 | 5,2E-04 |
| BASU050101 | Interactivity scale obtained from the contact matrix (Bastolla et al., 2005) | 0,70 | 5,5E-04 |
| LAWE840101 | Transfer free energy, CHP/water (Lawson et al., 1984) | 0,70 | 5,8E-04 |

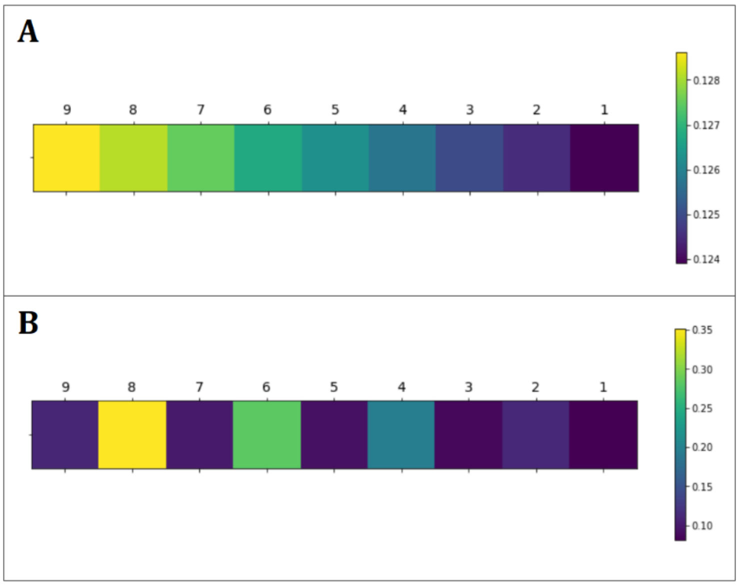


**Figure S1**. Graphical representation of obtained attention weights for proof-of-concept models 1 (A) and 2 (B).

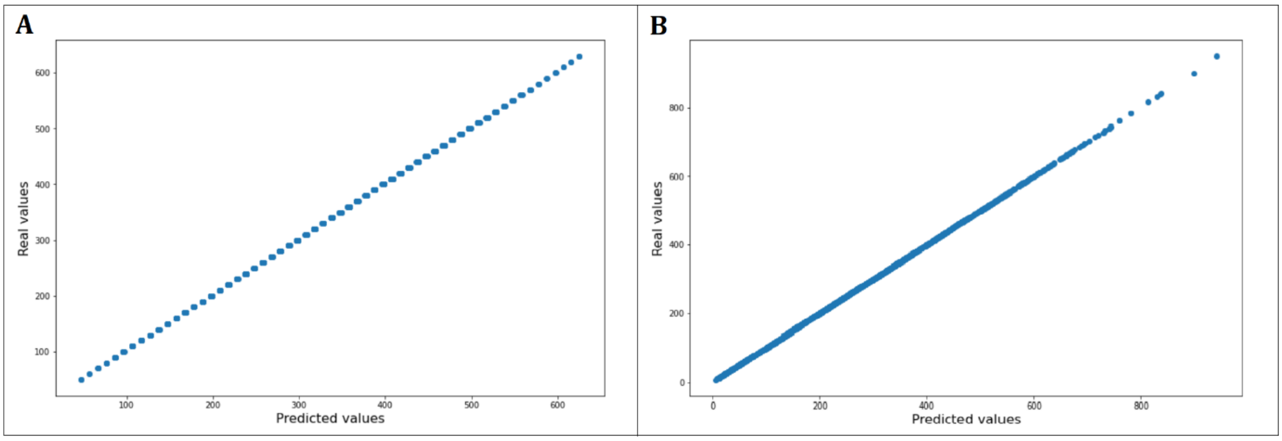


**Figure S2**. Graphical representation of the real versus predicted output values for proof-of-concept models 1 (A) and 2 (B).

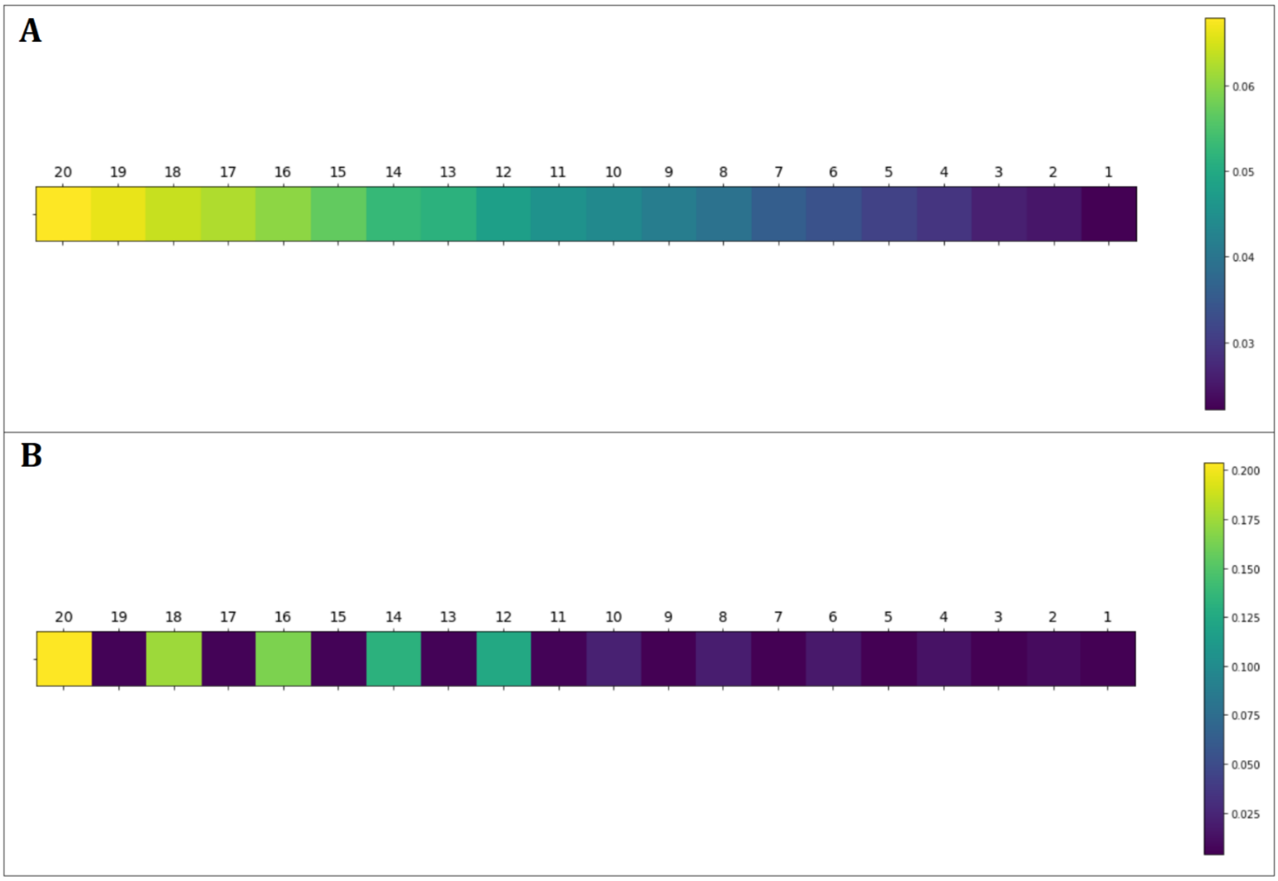


**Figure S3**. Graphical representation of attention weights for proof-of-concept models 3 (A) and 4 (B).

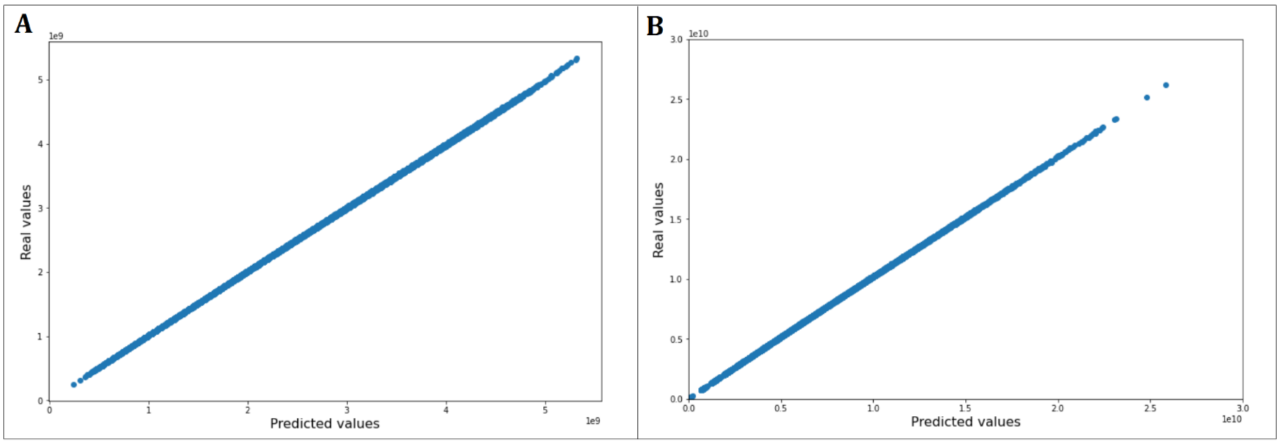


**Figure S4**. Graphical representation of the real versus predicted output values for proof-of-concept models 3 (A) and 4 (B).

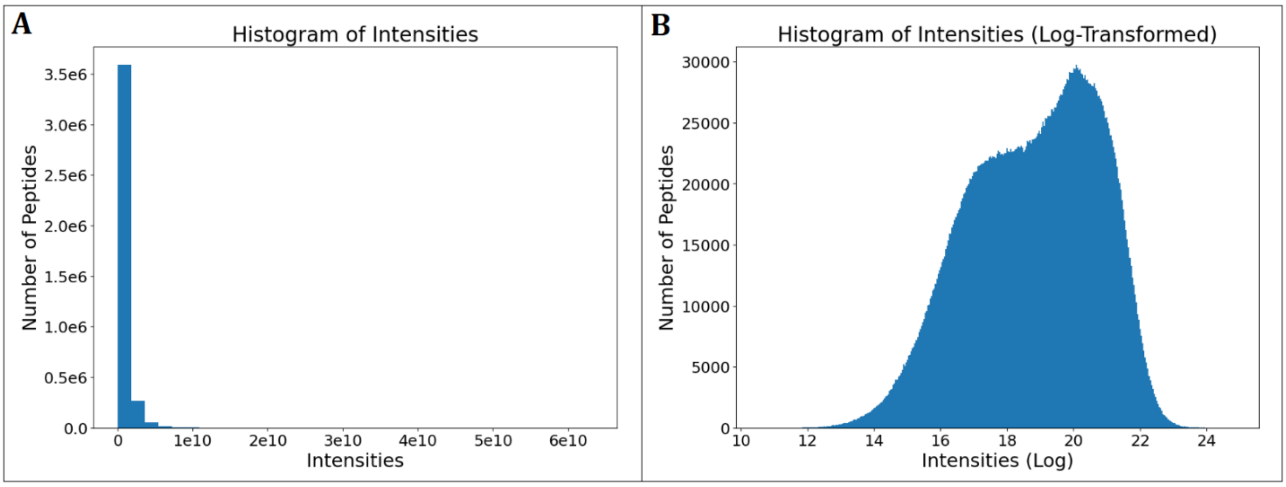


**Figure S5.** Distribution of the unfiltered data. A, Histogram of intensity distributions showing raw MS1 intensities (Bin size ≈ 1.8e9). B, Histogram of log-transformed MS1 intensities (Bin size ≈ 0.038).

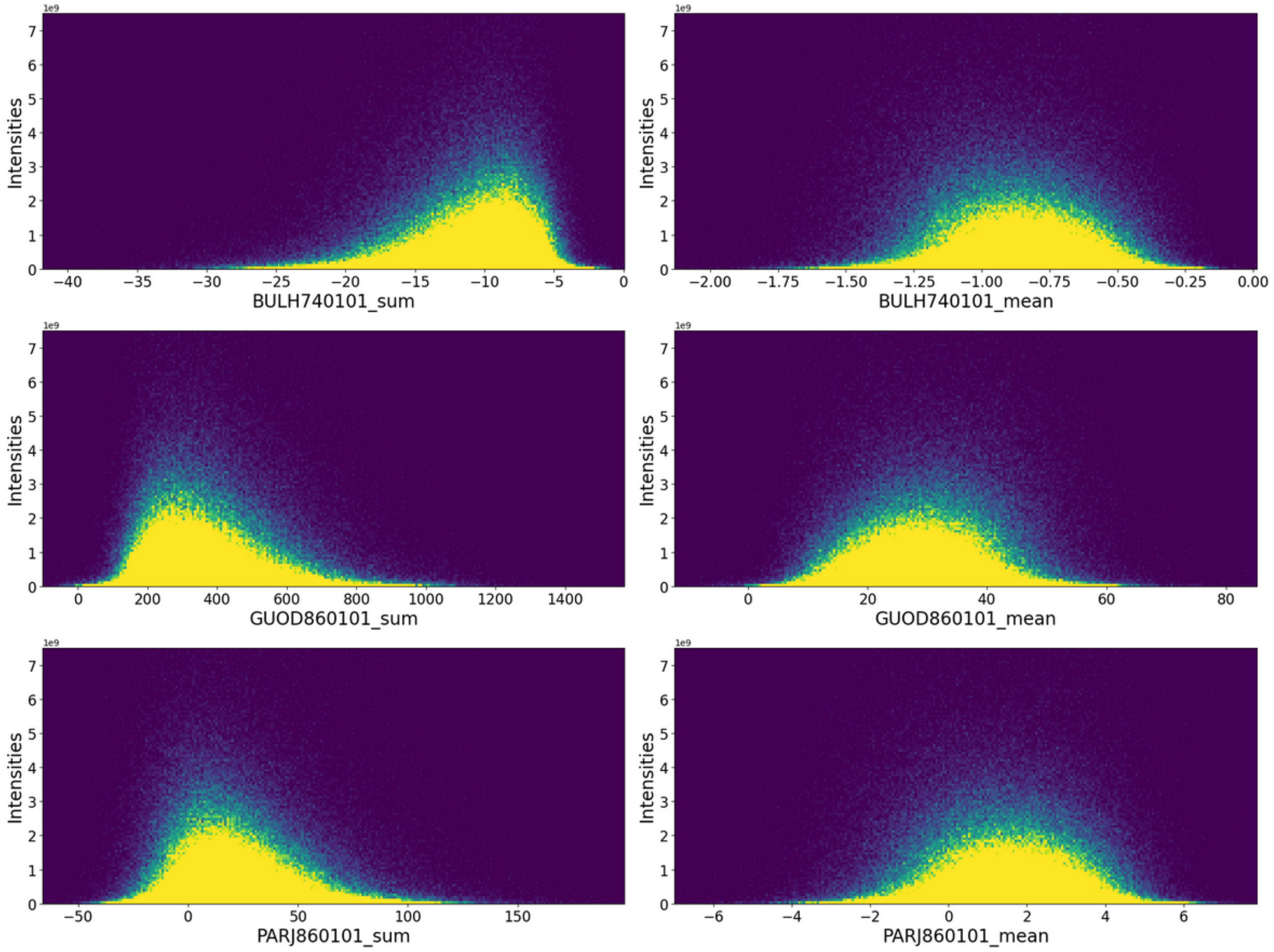


**Figure S6.** 2D histograms of MS1 intensities versus sum (left) and mean (right) computed hydrophobic properties of peptides based on AAindex indices BULH740101 (top), GUOD860101 (middle), and PARJ860101 (bottom) for the filtered dataset. The color coding indicates point density going from 0 (dark blue) to 40 (yellow). There are 250 bins in each axis, and the Y axis has been set a maximum intensity value of 7.5e9.

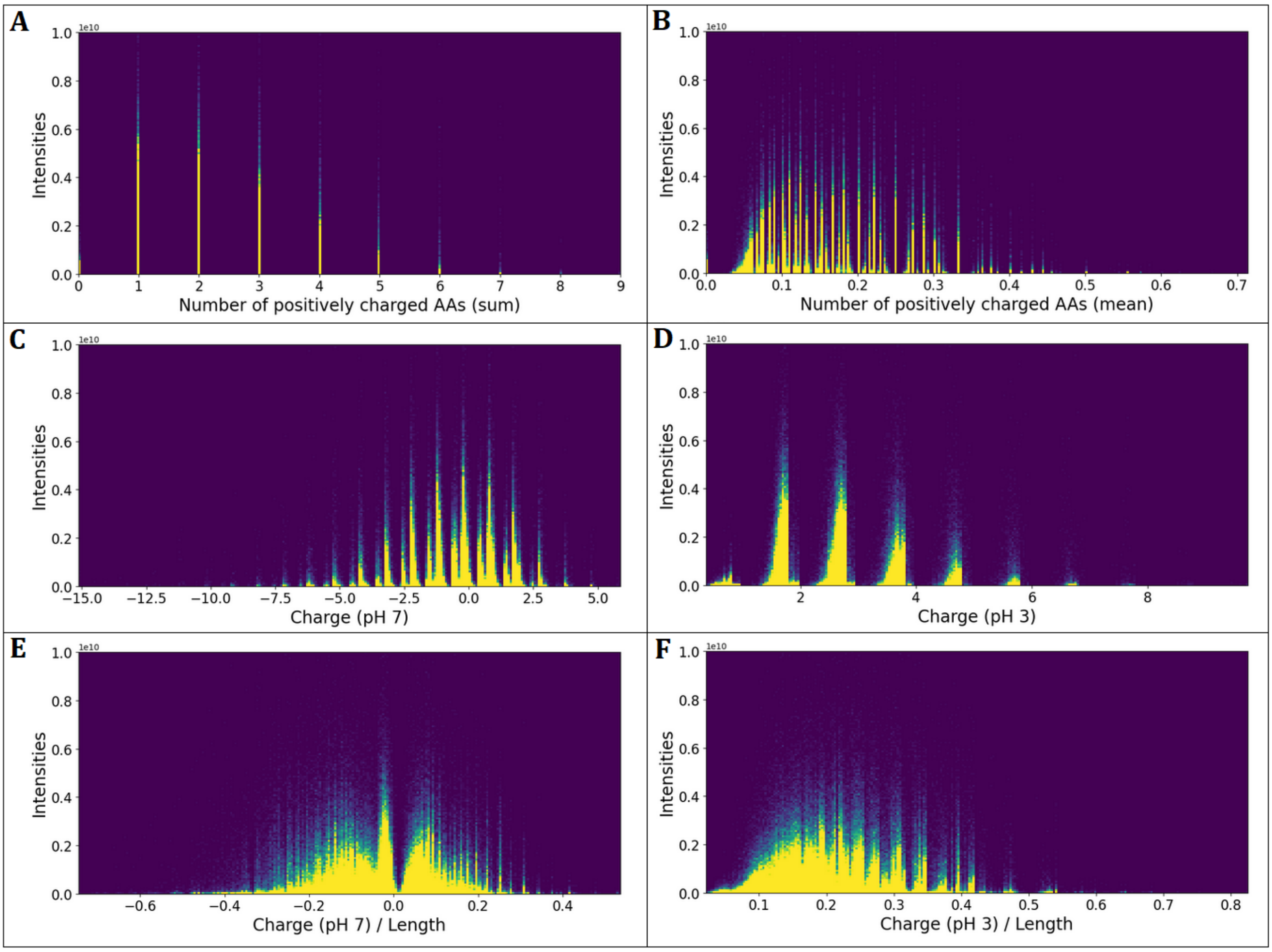


**Figure S7.** 2D histograms of MS1 intensities versus of computed charge for the filtered dataset. A, Sum of positively charged AAs. B, Mean of positively charged AAs. C, Charge at pH 7. D, Charge at pH 3. E, Charge at pH 7 (normalized by peptide length). F, Charge at pH 3 (normalized by peptide length). The color coding indicates point density going from 0 (dark blue) to 40 (yellow). There are 250 bins in each axis, and the Y axis has been set a maximum intensity value of 1.0e10.

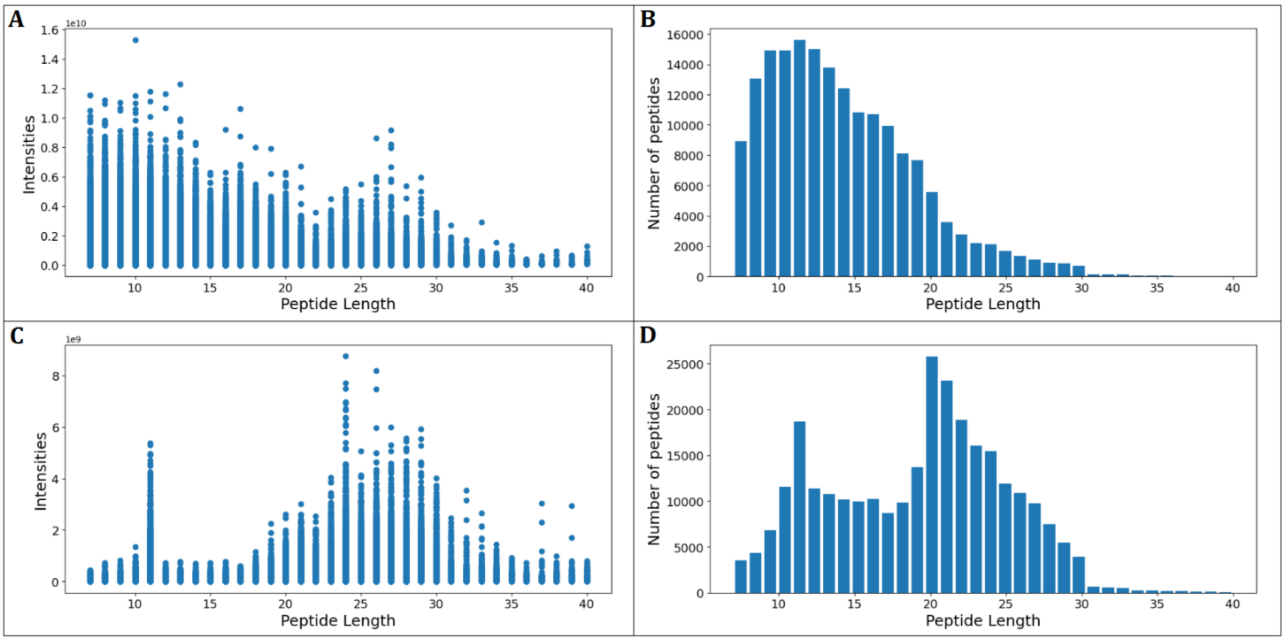


**Figure S8**. Graphical Representation of peptides for the filtered dataset and peptides identified with MaxQuant semi-specific search mode. A, Scatter plot of peptide length versus intensity in filtered dataset. B, Histogram of peptides length in filtered dataset. C, Scatter plot of peptide length versus intensity with MaxQuant semi-specific setting. D, Histogram of peptides length with MaxQuant semi-specific setting.

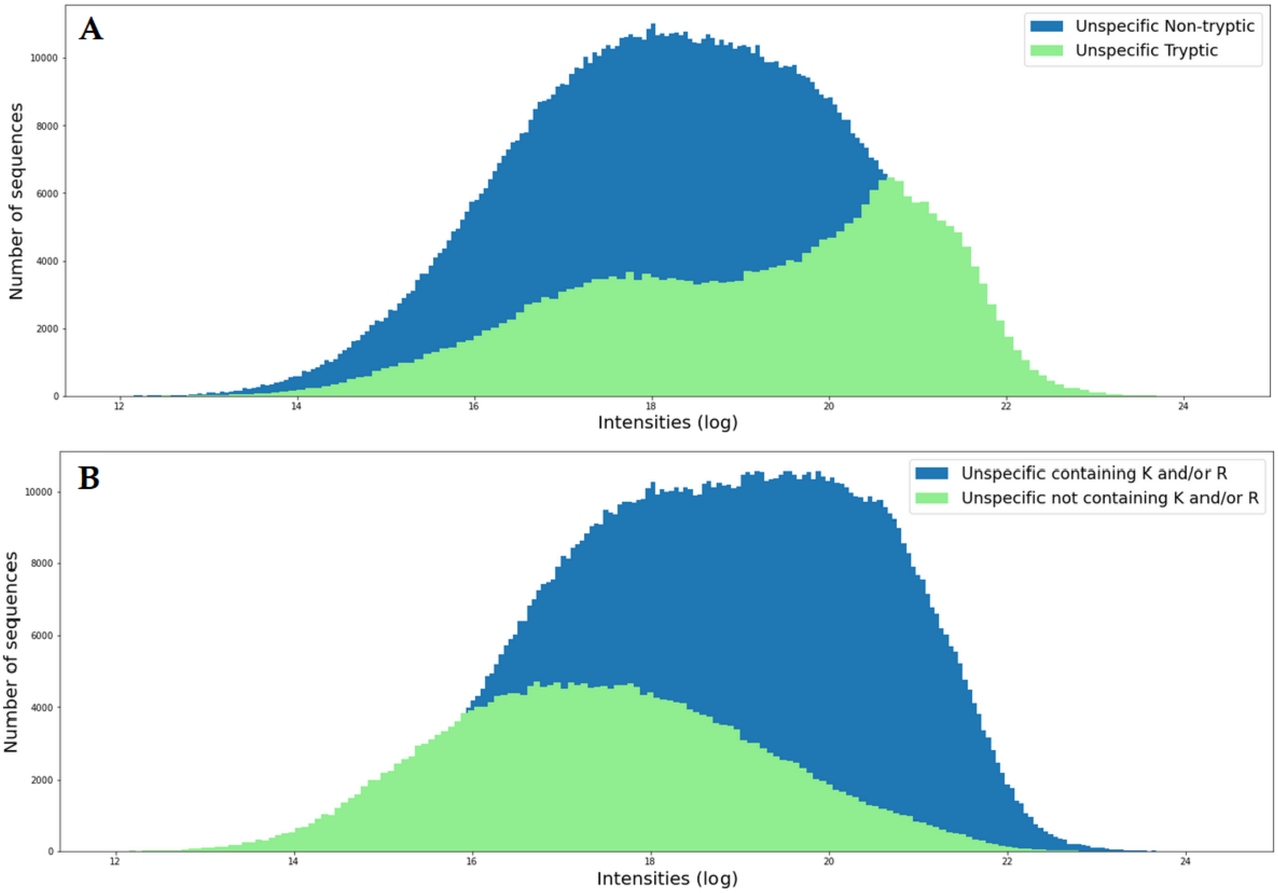


**Figure S9**. Histograms of peptides (Unspecific) and their log transform intensities. A, Histogram with all tryptic and non- tryptic peptides identified using unspecific search mode. B, Histogram of all peptides with and without K and/or R anywhere in the sequence, identified using unspecific search mode.

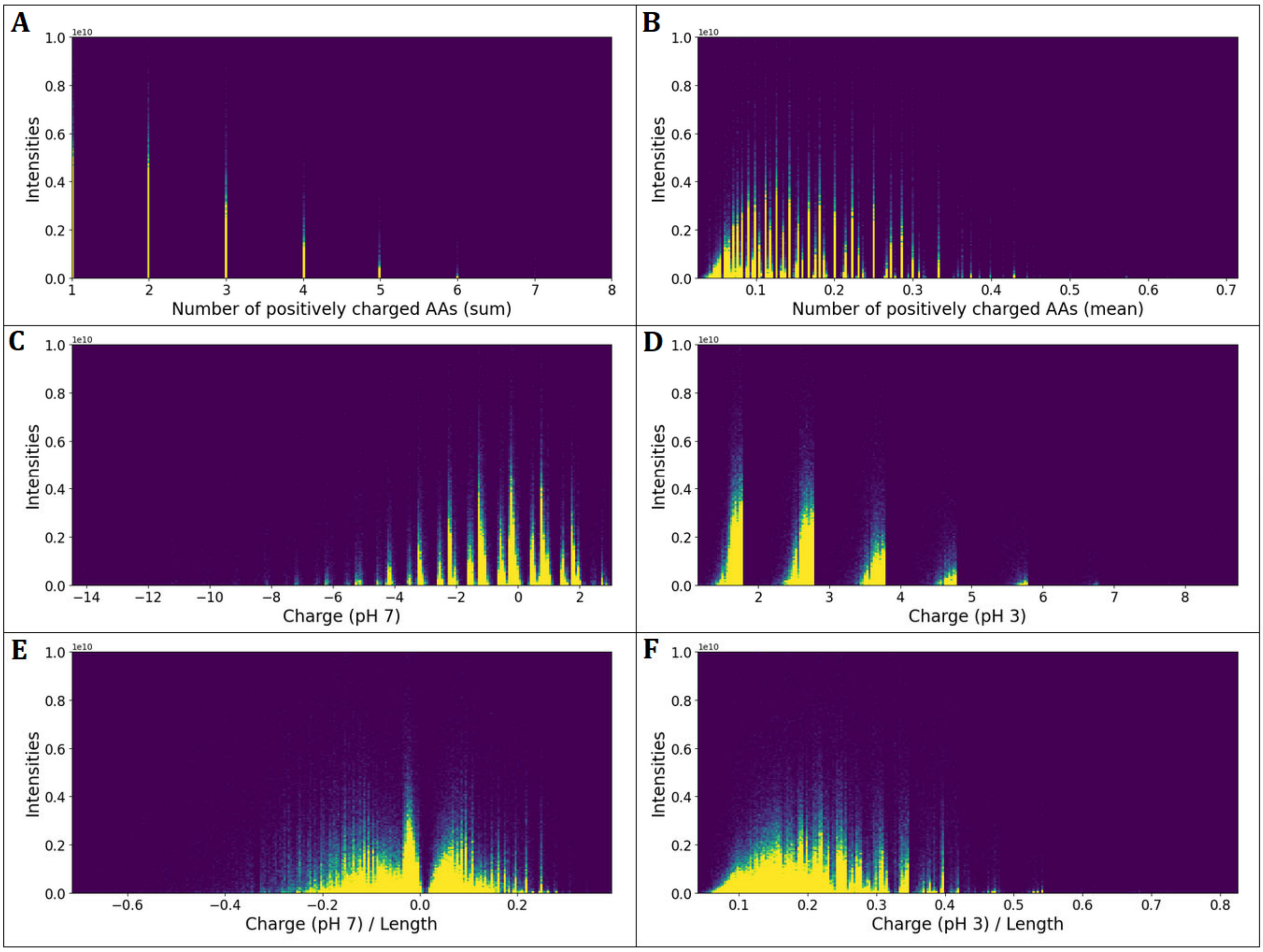


**Figure S10.** 2D histograms of MS1 intensities versus of computed charge for individual peptides in the specific/tryptic data subset. A, Sum of positively charged AAs. B, Mean of positively charged AAs. C, Charge at pH 7. D, Charge at pH 3. E, Charge at pH 7 over peptide length. F, Charge at pH 3 over peptide length. The color coding indicates point density going from 0 (dark blue) to 40 (yellow). There are 250 bins in each axis, and the Y axis has been set a maximum intensity value of 1.0e10.

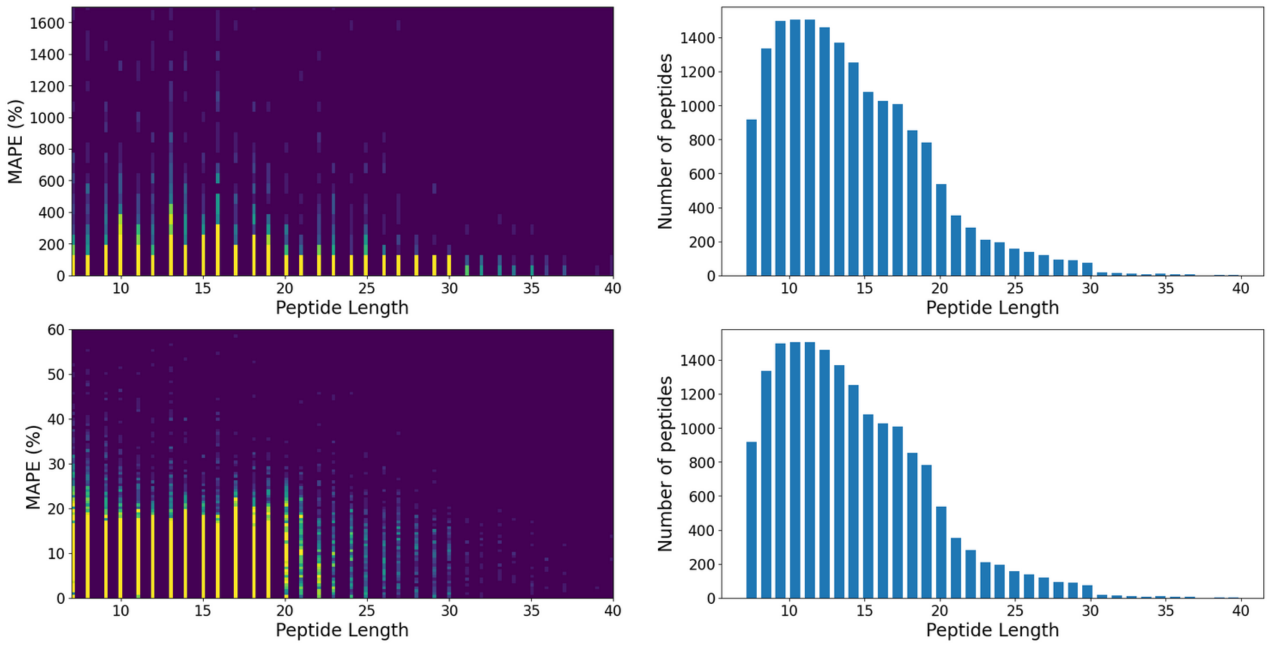


**Figure S11.** Distribution of MAPE for predicted MS1 intensities as a function of peptide length for the specific/tryptic test dataset. Left: 2D histograms of MAPE for predicted MS1 intensities prediction versus peptide length in real scale (top) and log-transformed (bottom). The color coding indicates point density going from 0 (dark blue) to 15 (yellow). There are 150 bins in each axis, and the Y axis has been set a maximum intensity value of 1700 (top) and 60 (bottom). Right: Histograms of peptide length (top and bottom are identical).
